# Synaptic vesicle undocking induces low frequency depression

**DOI:** 10.64898/2026.01.15.699619

**Authors:** Melissa Silva, Federico F. Trigo, Isabel Llano, Alain Marty

## Abstract

Synaptic depression is often interpreted as reflecting depletion of the readily releasable pool (RRP) following exocytosis. Such a mechanism predicts little or no depression at low stimulation frequency, as RRP replenishment should then offset the loss of vesicles by exocytosis. Nevertheless, in several types of mammalian central synapses, repetitive presynaptic stimulation at low frequency (< 5 Hz) elicits synaptic depression (low frequency depression, or LFD). In the present work we count the number of synaptic vesicles released at individual active zones to study the RRP and its replenishment during LFD. Contrary to depletion models of synaptic depression, we find that LFD does not depend on previous SV consumption. We find that LFD displays a long recovery time course (tens of seconds) when challenged by isolated stimulations but is immediately reversed by a high frequency train. We suggest that LFD results from undocking, a shift between two classes of synaptic vesicles organized sequentially inside the RRP (replacement vs. docked vesicles) in favor of the upstream (replacement) state. While undocking is apparent hundreds of milliseconds after a stimulation, calcium dependent docking takes only a couple of milliseconds, explaining the fast LFD recovery when stimulating at high frequency. Consistent with the undocking model, we find that double presynaptic stimulations alleviate LFD as they favor vesicular docking and RRP replenishment. Finally, we expand our model to explain how stimulation frequency shapes short-term synaptic depression, changing from depression at low frequency to a facilitation-depression sequence at medium or high frequency trains.

## Introduction

Short term synaptic plasticity (STP) governs changes in the amplitude of synaptic signals following recent neuronal activity. These changes are often rapid and large, and they potently shape information transfer across synapses. Two main components of STP are synaptic facilitation and synaptic depression (Zucker and Regehr, 2002).

Synaptic depression occurs after a burst of high frequency presynaptic firing and associated synaptic release (Betz, 1970; Castillo and Katz, 1954; Charlton et al., 1982). This link with previous presynaptic activity led to the view that synaptic depression is largely due to a decrease in the size of the readily releasable pool (RRP) of synaptic vesicles (SVs), following the loss of SVs by exocytosis (Kavalali, 2006; Regehr, 2012; Zucker and Regehr, 2002). In addition, a number of other mechanisms including desensitization of postsynaptic receptors (Rossi et al., 1995; Trussell et al., 1993), activation of presynaptic inhibitory receptors (Redman and Silinsky, 1994), decreased voltage-dependent calcium entry (Xu and Wu, 2005), decreased SV release probability (Wölfel et al., 2007; Wu and Borst, 1999), heterogeneous release probabilities among release sites (Neher, 2015; Trommershäuser et al., 2003), and non-linear presynaptic calcium buffers (Bolshakov et al., 2019), contribute to synaptic depression in various proportions depending on the preparation.

In more recent years, several studies have uncovered a form of synaptic depression that may differ from the RRP depletion pattern. In invertebrate synapses (*Aplysia* (Doussau et al., 2010); crayfish (Silverman-Gavrila et al., 2005)), as well in mammalian central synapses (hippocampal CA3 to CA1 (Abrahamsson et al., 2007) and CA3-CA3 (Saviane et al., 2002) synapses, climbing fiber to Purkinje cell synapses (Rudolph et al., 2011); parallel fiber to Purkinje cell synapses (Doussau et al., 2017); calyx of Held (Lin et al., 2022; Müller et al., 2010)), repetitive stimulation at low frequency (0.1 Hz to 5 Hz) induces synaptic depression. This form of synaptic depression (low frequency depression, or LFD) does not require a previous period of intense presynaptic activity, so that it is unclear whether it can be explained by the classical RRP depletion hypothesis, or by any of the alternative mechanisms mentioned above.

Recent studies suggest that the RRP is made up of a heterogeneous group of SVs, and that the relevant parameter for synaptic plasticity is not the RRP size as such, but rather the proportion between two components of the RRP (primed vs. unprimed or superprimed SVs; loosely vs. tightly docked SVs; or docked vs. replacement SVs) (Blanchard et al., 2020; Doussau et al., 2017; Eshra et al., 2021; Fukaya et al., 2023; Kim et al., 2024; Kobbersmed et al., 2020; Koppensteiner et al., 2022; Lin et al., 2022, 2025; Miki et al., 2020, 2016; Taschenberger et al., 2016). These results open new perspectives for the interpretation of STP (Neher, 2023; Neher and Brose, 2018; Pulido and Marty, 2018; Schmidt, 2019; Silva et al., 2021).

Electron microscopy studies of hippocampal synapses in culture showed that, following a single presynaptic action potential (AP) stimulation, the distance between SVs and the active zone (AZ) membrane varies as a function of time in a complex manner (Kusick et al., 2020; Ogunmowo et al., 2025). Following the immediate loss of SVs by exocytosis, the number of SVs directly in contact with the AZ membrane rapidly increases (within ∼15 ms), reflecting the movement of SVs from a cytosolic location to the membrane (‘calcium-dependent docking’). Within ∼100 ms, SVs move back into the cytosol (‘calcium-dependent undocking’). In the following several seconds, SVs partially move forward again to the AZ membrane as the system returns to its basal state. While SV docking has been associated with facilitation (Miki et al., 2016; Schmidt, 2019; Silva et al., 2021), the functional role of subsequent undocking has remained unclear. Here we explore the possibility that undocking could underly LFD.

Testing the undocking hypothesis of LFD necessitates a functional assay of various subclasses of RRP SVs, a notoriously difficult task (Neher, 2015). Our laboratory has developed a method to monitor the release of individual SVs at single AZ synapses between parallel fibers (PFs) and molecular layer interneurons (MLIs) in cerebellar slices (‘simple synapse recording’ (Malagon et al., 2016)). Simple synapse recording has provided models of synaptic facilitation and of high frequency depression based on changes in occupancy of docking sites and associated SV groups inside and outside the RRP (Miki et al., 2016; Tanaka et al., 2021; Tran et al., 2022). In the present work, we suggest that LFD reflects a specific SV re-equilibration inside the RRP, as well as an inhibition of SV entry into the RRP. We further propose a general model of synaptic depression which explains differences between low and high stimulation frequencies.

## Results

### Biphasic dependence of paired-pulse ratio on inter-stimulus interval

PF-MLI synapses are considered facilitating (Atluri and Regehr, 1998). In the present work, we ask whether single PF-MLI synapses may turn depressing at low frequency. Using simple synapse recordings at PF-MLI junctions, short trains of 4 presynaptic APs were applied at various inter-stimulus intervals (ISIs) in a scrambled order. Each train was separated from the next by a 10 s-long time interval, considered long enough to reset the system (Tran et al., 2023) (**Fig. 1A**). The resulting EPSCs were deconvolved using the mean quantal EPSC as kernel to yield the numbers of released SVs per AP (s_i_, where i stands for AP number) (Malagon et al., 2016). Results of an example experiment are shown in **Fig. 1 B-C**. **Fig. 1C, top** shows average EPSC traces for the first AP (average s_1_) and for the second AP at various ISI values. **Fig. 1C, bottom** shows, for each ISI, the ratio of mean released SV numbers for AP #2 over the corresponding mean for AP #1 (paired pulse ratio; PPR: blue dots; results for AP #3-4 are shown in **Supplementary Fig. 1**). In this experiment, the PPR defined in this manner displayed values > 1 for ISIs ranging from 10 to 100 ms, and values < 1 for ISIs ranging from 200 to 1600 ms. In group results, the PPR was consistently > 1 for ISI values of 100 ms or less, and < 1 for ISI values of 800 or 1600 ms (**Fig. 1D**). A biexponential fit to the data (red curve in **Fig. 1D**) intersected the PPR = 1 line for an ISI near 300 ms. These results show that the synapse changes from facilitating to depressing depending on the ISI. They differ from the single facilitation component previously reported for this synapse (Atluri and Regehr, 1998). This discrepancy presumably stems from differences in experimental conditions (room temperature, stimulation of multiple presynaptic PFs and 2 mM external Ca^2+^ concentration in the previous work, vs. near-physiological temperature, single presynaptic stimulation and 3 mM external Ca^2+^ here; see below). Here, the PPR is the sum of a fast facilitation component and of a slow depression component (PPR = 1 + A_fast_ exp(-t/τ_fast_) - A_slow_ exp (-t/ τ_slow_), with A_fast_ = 0.94, τ_fast_ = 230 ms, A_slow_ = 0.30, τ_slow_ = 2 100 ms).

**Figure 1:**
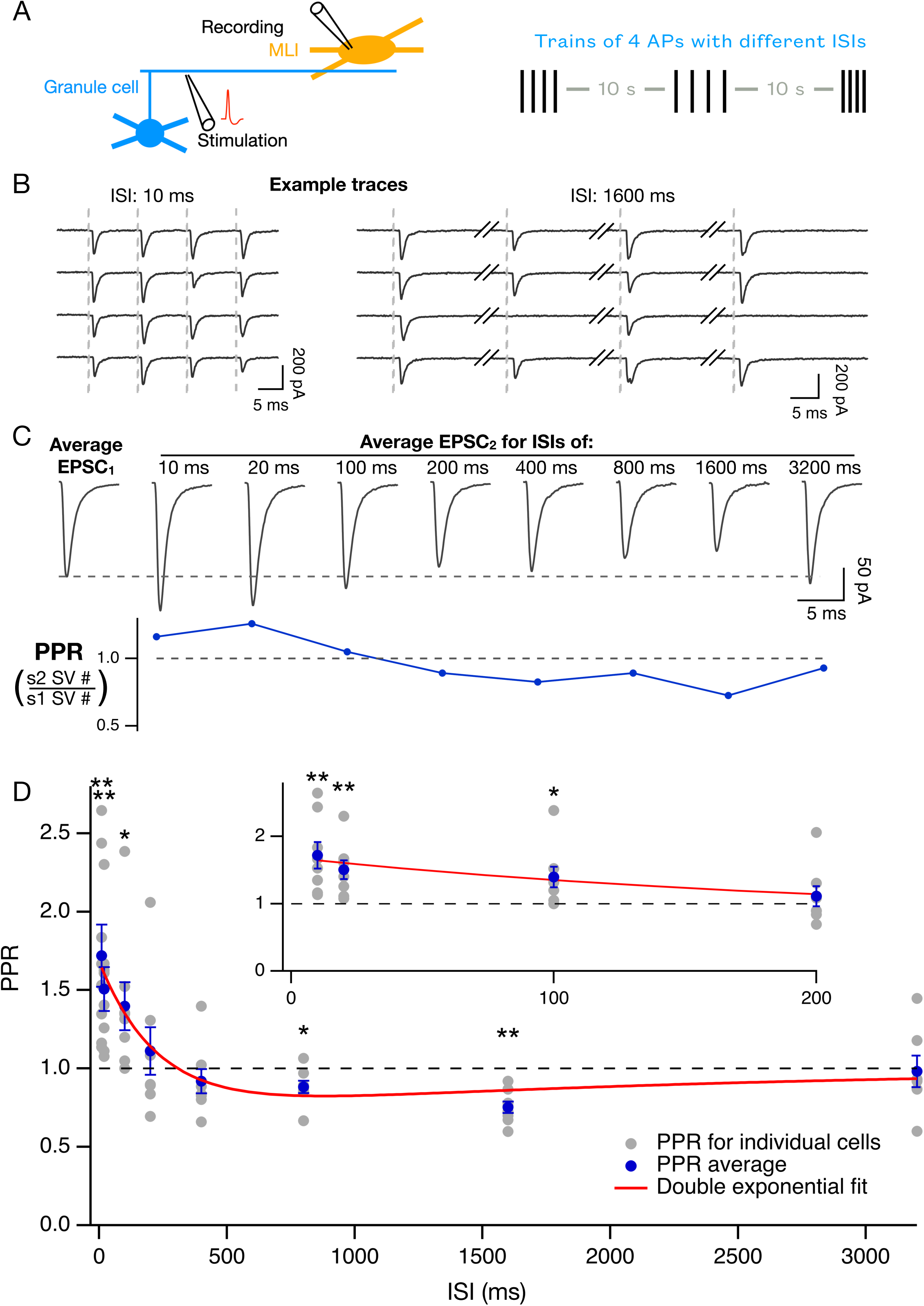
Dependence of paired-pulse ratio on inter-stimulus interval. **A**: Experimental protocol. To obtain simple synapse recordings, individual granule cell axons were stimulated with an extracellular pipette located in the granule cell layer, and EPSCs were recorded in a postsynaptic MLI. Presynaptic stimulations involved trains of 4 action potentials with various inter-stimulus intervals (ISIs), and with 10 s inter-train intervals. **B**: Example recording showing responses to trains using 10 ms ISIs (left) and 1 600 ms ISIs (right). Stimulation times indicated by dotted vertical lines. **C, Top**: Average EPSC in response to AP #1 (1st trace, labelled ’average EPSC1’) followed by average responses to AP #2 for ISIs varying from 10 ms to 3200 ms, from the same experiment (20 repetitions for each ISI). **C, Bottom**: A measure of the paired-pulse ratio, PPR, calculated from the ratio of the mean numbers of released SVs for AP #2 over the average release for AP #1 from the same data. Numbers of SVs released by individual presynaptic stimulations were determined by deconvolution, using the mean quantal EPSC as kernel. **D**: Plot of PPR as a function of ISI (same analysis as in **C, bottom:** grey dots, individual cells (means from > 20 repetitions for each cell); blue dots and associated error bars, means ± sem from n = 8 cells). The red curve is a double exponential fit to the data with A_fast_ = 0.94, τ_fast_ = 230 ms, A_slow_ = 0.30, τ_slow_ = 2100 ms. **Inset:** Blow-up of results for ISIs of 200 ms or less. * and ** indicate data points that differ from PPR = 1 with p < 0.05 and p < 0.01 respectively.

### Low frequency depression (LFD) can be modelled as reflecting a decrease in DS occupancy but no change in RS occupancy or in IP size

We previously developed a sequential 2-step model of docking at PF-MLI synapses (Miki et al., 2018, 2016; Tran et al., 2022) (**Fig. 2A**). In this model, called replacement site/docking site (RS/DS) model hereafter, each AZ contains a fixed number of docking units (DUs), with a mean value of 4 DUs per AZ (Miki et al., 2017). Each DU comprises two SV binding sites: a docking site (DS), which is directly attached to the AZ membrane, and an associated replacement site (RS). The RS can accommodate an SV regardless of whether the associated DS is occupied or not, meaning that a DU can simultaneously bind two SVs (Silva et al., 2024). The RRP is the sum of the SVs bound to all DUs (RS and DS) in the same AZ. High frequency AP trains induce a flow of SVs from an upstream intermediate pool (IP) to the RRP. Once in the RRP, SVs transit from the RS to the DS, and eventually undergo exocytosis (downward red arrows in **Fig. 2A**).

**Figure 2:**
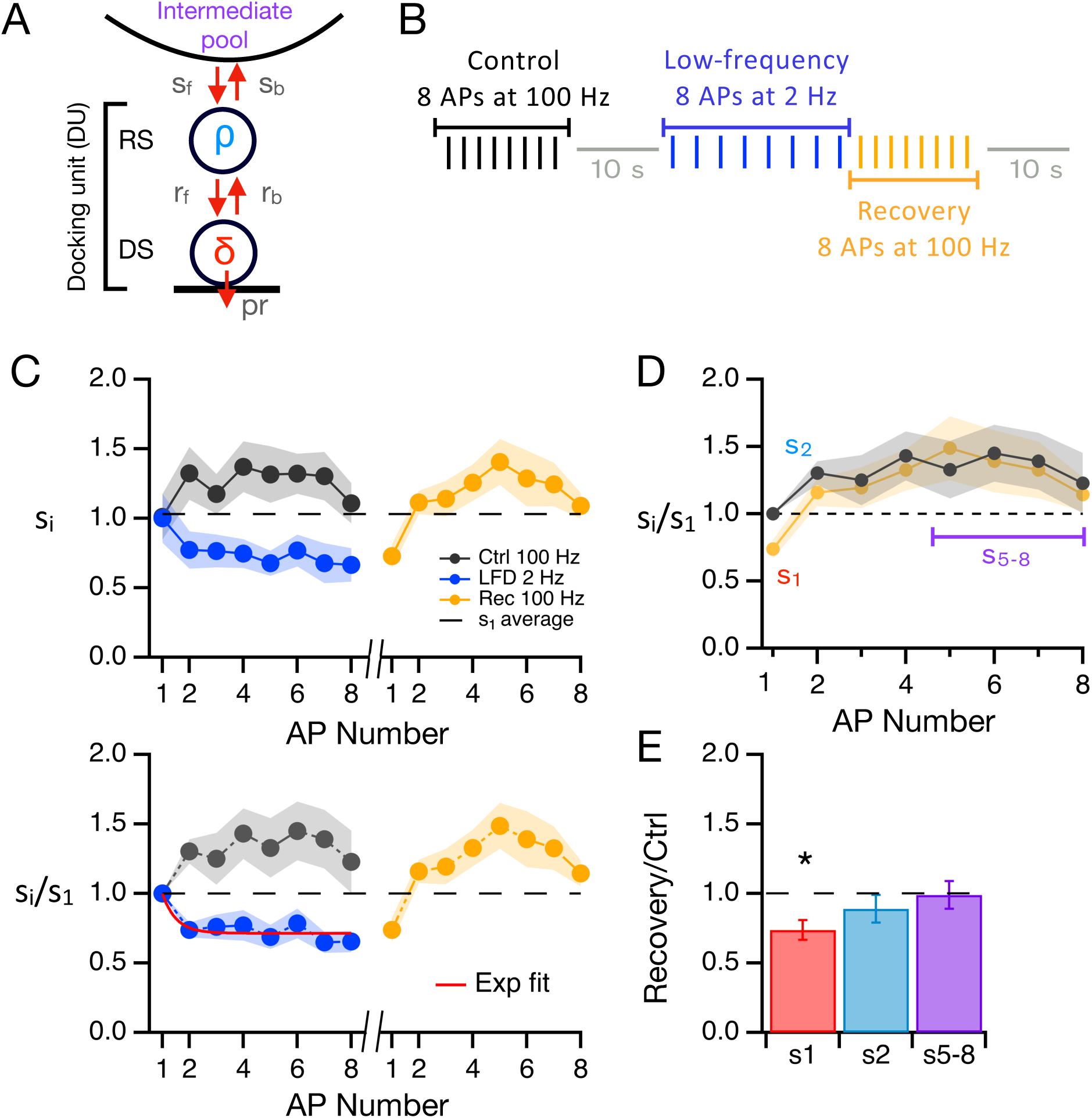
LFD reflects a decrease in docking site occupancy. **A**: Sequential 2-state docking model. SVs coming from the intermediate pool (IP) transit through a replacement site (RS, with occupancy ρ) and an associated docking site (DS, with occupancy δ) before release. Taken together, one RS and one DS constitute one docking unit (DU). s_f_, s_b_, r_f_ and r_b_ represent transition rates as indicated. p_r_ represents the release probability of a docked SV following an AP. **B**: Experimental protocol. To probe the state of SV pools in the synapse, trains of 8 APs at 100 Hz were applied either in isolation (control train), or after a depressing train of 8 APs at low stimulation frequency (recovery train). **C, Top**: Plots (means ± sem from n = 7 cells; > 20 repetitions for each cell) of s_i_ for the control train (black), for the low frequency train (2 Hz, blue), and for the recovery train (yellow). Dotted line: Average s_1_ value from control and low frequency trains. **C, Bottom**: Same, after normalization with respect to s_1_ value. Exponential fit with asymptotic value = 0.69 ± 0.05 and time constant = 0.87 AP # (= 435 ms; red curve). **D**: Superimposed s_i_ plots for control (black) and recovery (yellow). Comparison of the SV release between control and recovery trains allows us to evaluate the occupancy of the RRP and of the IP at the end of the low frequency train. SV numbers in response to the 1st (s_1_), 2nd (s_2_), and last (s_5-8_) APs respectively report changes in δ (red), in ρ (blue), and in the IP size (purple). **E**: Ratios of s_i_ values between recovery 8-AP train and control train. s_1_ is significantly reduced, indicating a decrease in δ, but neither s_2_ nor s_5-8_ are changed, indicating no change in ρ or in IP size.

The RS/DS model of **Fig. 2A** has been used to interpret phenomena of STP during and after high frequency AP trains. In response to stimulation at 100 or 200 Hz, the s_i_ plot displays an initial increase up to i = 2-4, reflecting synaptic facilitation, followed by a gradual decrease reflecting synaptic depression (Miki et al., 2016; Tran et al., 2022). According to the model, the DS occupancy, δ, increases after a stimulus over its basal value, following Ca^2+^-dependent movement of SVs from RS to DS. This explains the initial facilitation. Subsequent synaptic depression reflects a δ decrease following the gradual exhaustion of IP SVs and consequently, a decrease in the flow of SVs into the RRP. Recovery requires the refilling of the IP, and it occurs on a time scale of hundreds of ms.

To explore the mechanisms underlying STP as a function of AP frequency, we alternated 8-AP stimulations at 100 Hz and at 2 Hz. Immediately after the 2 Hz train, we repeated the 100 Hz train, before repeating the entire protocol again after a pause of 10 s, to allow for synapse recovery (**Fig. 2B**). According to our previous work (Tran et al., 2022), and within the framework of the RS/DS model, we can estimate the occupancy of the RRP and the intermediate pool (IP) during the low frequency stimulation by comparing the s_i_ curves of the 100 Hz trains before (control) and immediately after (recovery) the 2 Hz train. Under the hypothesis that the release probability of docked vesicles (p_r_) remains constant throughout the AP train, changes in δ, in RS occupancy (ρ) and in IP size are reflected by changes in the number of released SVs for AP #1 (s_1_), for AP #2 (s_2_), and for the average of AP #5-8 (s_5-8_) respectively. This analysis was derived under conditions of elevated release probability. Therefore, in most of our experiments, we used an increased external Ca^2+^ concentration (3 mM) to facilitate the interpretation of the results in terms of changes in RRP and IP occupancy; results obtained under physiological external Ca^2+^ concentration conditions will be reported below.

The results of these experiments are shown in **Fig. 2C-E**. In keeping with the results of **Fig. 1**, when stimulating at 2 Hz in 3 mM external Ca^2+^ concentration, a clear synaptic depression was observed (PPR = 0.74 ± 0.05, paired one-tailed t-test, p = 0.0027, n = 7; blue traces in **Fig. 2C**; absolute SV numbers in **Fig. 2C top**, and values normalized to control s_1_ value in **Fig. 2C bottom**). The s_i_ curve for 2 Hz trains could be approximated to an exponential decay with an asymptote of 0.69 ± 0.05 (paired one-tailed t-test, p = 0.0032, n = 7) and a time constant of 0.87 ISIs (435 ms; red curve in **Fig. 2C bottom**). We then compared release during control and recovery trains to investigate the origin of the synaptic depression (**Fig. 2D**). The value of s_1_ at the onset of the recovery train replicated the stable low s_i_ values observed at the end of the low frequency train. It displayed a decrease with respect to the s_1_ value of control trains (to 70 % of this control; p = 0.02, one-tailed paired t-test; **Fig. 2C bottom**). As s_1_ is a proxy for δ under the present experimental conditions, this result suggests that LFD involves a decrease in δ, as previously suggested for high or moderate frequency depression (HFD) (Borges-Merjane et al., 2020; Lin et al., 2022; Miki et al., 2016; Tanaka et al., 2021). Strikingly, s_i_ values during the recovery train caught up with control values immediately after the first stimulation, so that neither s_2_ nor s_5-8_ values differed from those of control 100 Hz trains (s_2_ = 1.30 ± 0.09 for control and 1.16 ± 0.10 for recovery; p > 0.05; s_5-8_ = 5.05 ± 0.68 for control and 5.02 ± 0.63 for recovery; p > 0.05, one-tailed paired t-test; compare yellow and black s_i_ curves in **Fig. 2D**; summary graph in **Fig. 2E**). Within the framework of the RS/DS model, these results suggest that neither ρ nor the IP size were decreased at the end of the low frequency train. Taken together, the results suggest similarities and differences between the mechanisms of synaptic depression at high frequency, as studied before, and at low frequency, as studied here. In both cases, the RS/DS model suggests that synaptic depression is accompanied by a decrease in δ. However, while ρ and IP pool size accompany the δ decrease during synaptic depression at high stimulation frequency (Tanaka et al., 2021; Tran et al., 2022), only δ is changed during LFD.

### LFD as a function of stimulation frequency and external calcium concentration

Having documented LFD at 2 Hz stimulation frequency and in 3 mM external Ca^2+^ concentration as shown in **Fig. 2**, we next examined the results of similar experiments when changing the stimulation frequency to 1 Hz, or when reducing the external Ca^2+^ concentration to near physiological concentration (1.5 mM). When stimulating at 1 Hz, LFD was obtained (**Supplementary Fig. 2A**, blue triangles; fractional response at steady state: 0.79 ± 0.08, paired one-tailed t-test, p = 0.0037, n = 5). As was the case with 2 Hz stimulations, the s_i_ curve recovered immediately after the first stimulus during the recovery train (yellow curves in **Supplementary Fig. 2A)**. Next, we examined synaptic responses at 2 Hz in 1.5 mM external calcium, finding no depression (**Supplementary Fig. 2B**). At this calcium concentration, synapses facilitate more than at 3 mM (compare control black normalized traces in **Fig. 2C bottom** in 3mM Ca^2+^, and **Supplementary Fig. 2B-C right** in 1.5 mM Ca^2+^), potentially causing a longer-lived facilitation that would mask the depression. To test this theory, we performed the same protocol at 0.5 Hz in 1.5 mM Ca^2+^. While the PPR was not significantly < 1 (PPR = 0.83 ± 0.11, n = 6; p = 0.21, paired two-tailed t-test), the rest of the train led to a reduced number of released SVs compared to control s_1_ (fractional response at steady state: 0.69 ± 0.05, n = 6; p = 0.00058, paired one-tailed t-test), indicating LFD. An exponential fit to the 0.5 Hz data (red curve in **Supplementary Fig. 2C, right**) indicated a τ of 1.8 stimuli (3.6 s), slower than that observed for higher calcium concentrations at 2 Hz. The recovery curve (yellow, **Supplementary Fig. 2C**) showed a full recovery to control values once high frequency stimulation was resumed.

In conclusion, LFD can occur at 0.5 Hz, 1 Hz or 0.5 Hz; it occurs at near-physiological external Ca^2+^ concentration (1.5 mM), but a lower frequency is required compared to 3 mM Ca^2+^ (0.5 Hz instead of 2 Hz). These experiments further confirm that LFD is immediately reversed upon stimulation with high frequency.

### Modelling the dependence of PPR on ISI

The facilitation/depression curve of **Fig. 1** matches the docking/undocking sequence reported in flash-and-freeze experiments (Kusick et al., 2020). Therefore, we next asked whether the RS/DS model (**Fig. 2A**) could account for the dependence of the PPR on ISI on the basis of δ changes (constant p_r_ hypothesis). The sequence of events following an AP is shown in **Fig. 3**. Immediately after an AP, docked SVs were released with a probability p_r_, resulting in a drop in δ (grey area, **Fig. 3A**). We modelled subsequent SV movements based on transition rates r_b_, r_f_, s_b_, s_f_ (**Fig. 2A**; (Lin et al., 2022; Miki et al., 2018)) with appropriate time-dependent changes (**Supplementary Fig. 3**, **Table 1**, and **Materials and Methods**). This revealed a time window comprised between 10 and 200 ms when δ (red curve) was larger than its resting value of 0.6 (dotted red line, **Fig. 3A**), while ρ (blue curve) was lower than its resting value of 0.9 (dotted blue line, **Fig. 3A**). This time period corresponds with the facilitating part of the PPR vs. ISI plot (**Fig 3B**). Due to the low ρ value, SV entry into the RRP (IP to RS transition) was facilitated. We call this the ‘RS gate open’ configuration (green shade in **Fig. 3A**). By contrast, during the depressing part of the curve (**Fig. 3B**, ISI values comprised between 500 ms and 10 s), δ was smaller than its resting value while ρ recovered near its resting value, and the ‘RS gate’ was closed (an SV cannot move into the RS if it is already occupied; red shade in **Fig. 3A**). The predicted PPR plot (red curve in **Fig. 3B**) is proportional to the δ curve of **Fig. 3A** under the assumption of constant p_r_. It displays a good fit to the data (black dots), showing that the model can account for the facilitation/depression sequence of **Fig. 1** based on changes in DS occupancy.

**Figure 3:**
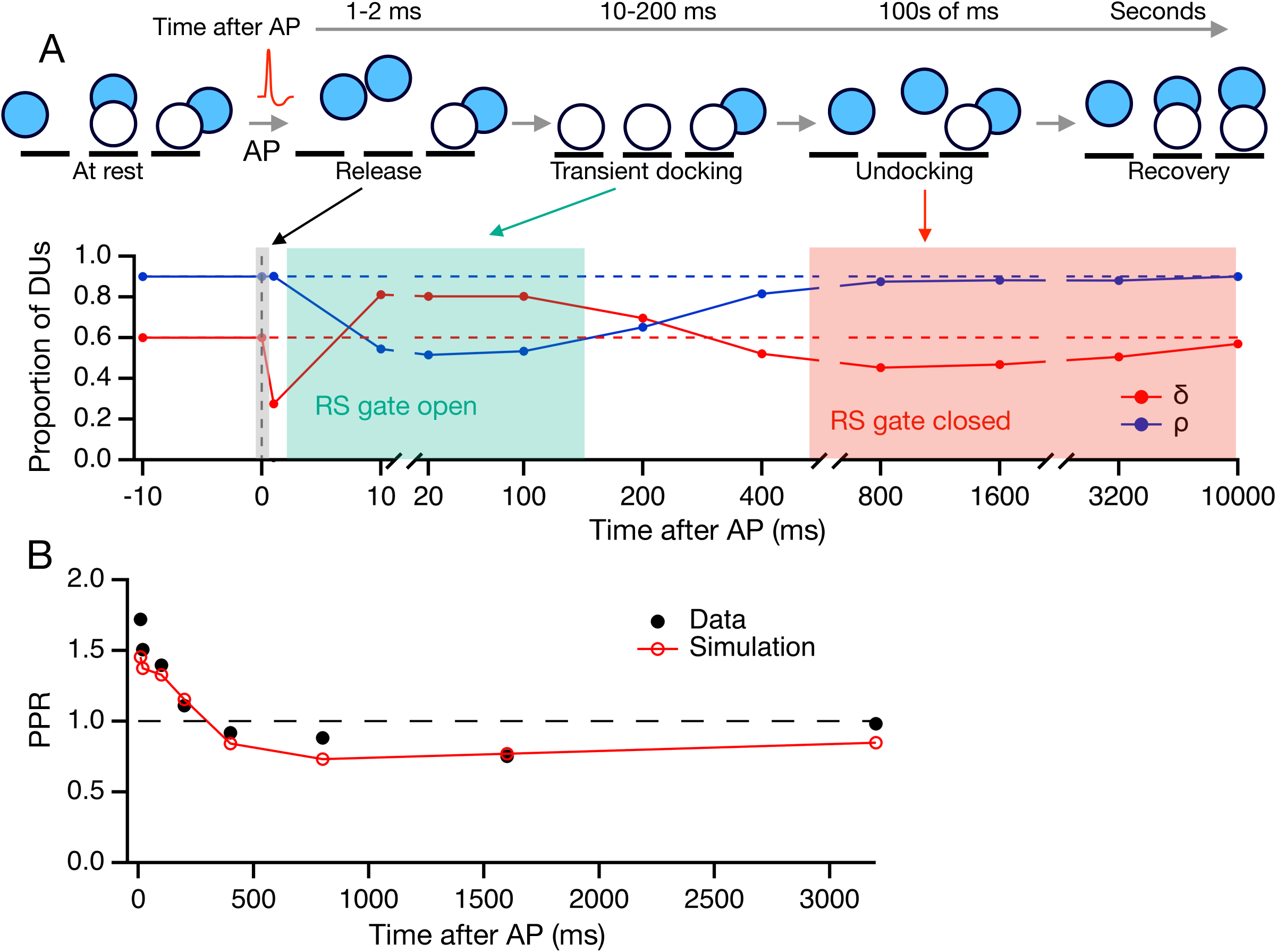
Simulations of s, δ and ρ curves after an AP stimulation. **A, up**: Model depiction of an AZ with 3 DUs at various time periods before and after an AP stimulation. After release, SVs transition from RS (blue; some of these SVs are placed sidewise as the 2-step model cannot specify the exact location of the RS) to DS (white) within tens of ms (transient docking), then after hundreds of ms they undock, before eventually returning to their basal state. **A, bottom**: Simulated time course of δ (red curve) and ρ (blue curve) before and after a presynaptic AP (at time 0). Resting δ and ρ values indicated by dotted lines. During transient docking, δ is high and ρ is low, so that RRP replenishment is allowed (RS gate open, green shade). During undocking, δ is low and ρ is high, so that RRP replenishment is blocked (RS gate closed, red shade). **B:** PPR curve as a function of ISI (black dots, from Fig. 1) together with simulation results (red curve).

**Table 1.**
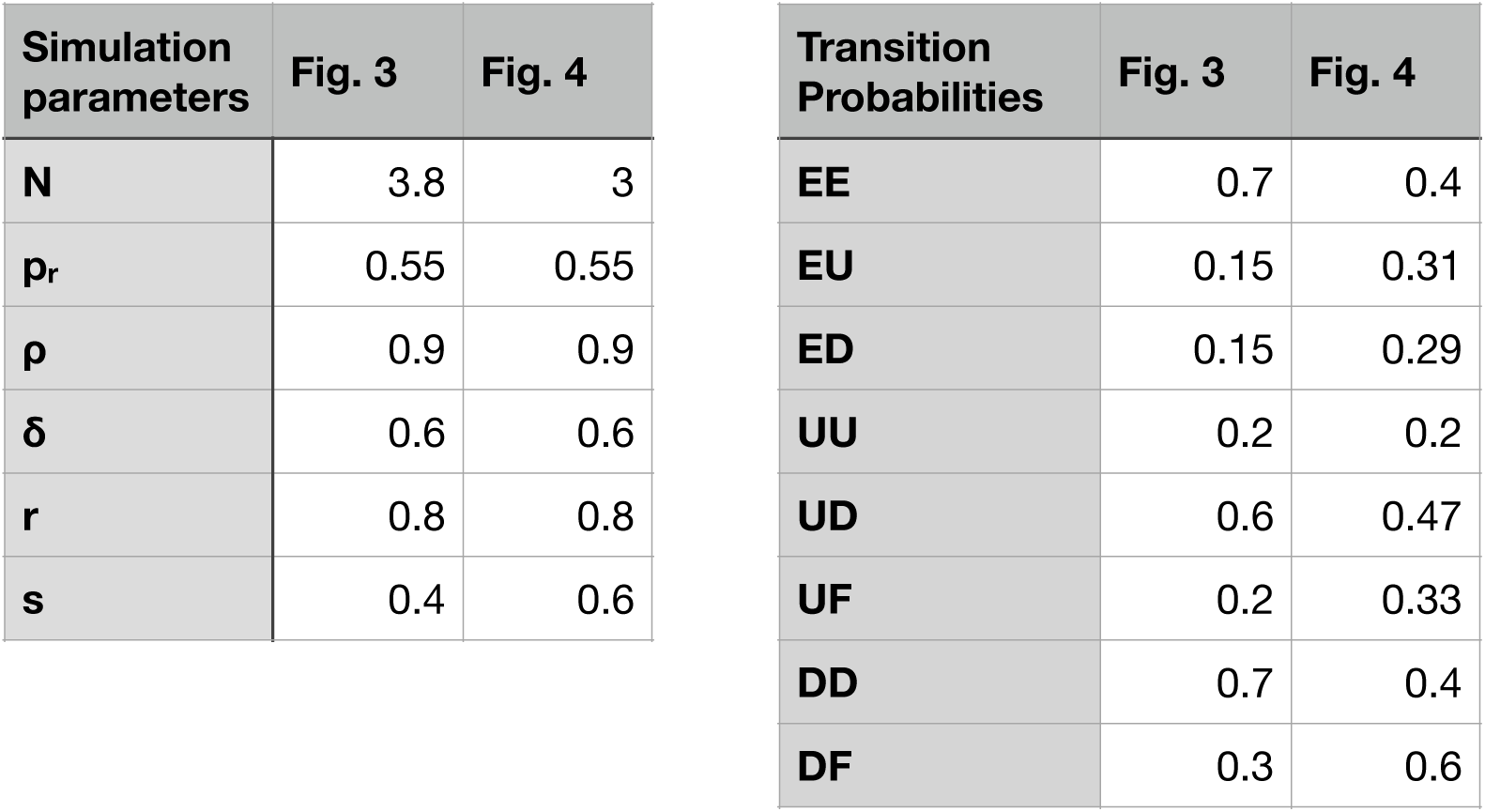

### Modelling synaptic output in response to AP trains at high or low frequency

Next, we examined whether the same model could account for the results of **Fig. 2** (LFD for 8-AP trains and following recovery). To this end, we made concatenations of the previous model over consecutive time segments (**Materials and Methods**). Data and simulations of 8-AP trains at high frequency (**Fig. 4A**), low frequency (**Fig. 4B**), and recovery trains at high frequency (**Fig. 4C**) illustrate the consequences of the time-dependent changes of DU occupancy shown in **Fig. 3**. Upper panels (**Fig. 4A, B** and **C**) show s_i_ plots together with simulation results, indicating that the model can reproduce the main features of **Fig. 2C** data. Lower panels (**Fig. 4A, B and C**) show plots of δ and ρ as a function of AP number. At high stimulation frequency (**Fig. 4A**), δ increases and remains high throughout an 8-AP train, explaining facilitation (red curve). Meanwhile, ρ decreases below its resting level (blue curve). Because SV movement from the IP into the RRP requires a free RS, this ρ decrease facilitates RRP replenishment. In effect, the RS gate opens at AP #2 and remains open throughout the 8AP train (**Fig. 4A, bottom; Fig. 4D, green shade**). At low stimulation frequency, a totally different pattern of changes is observed (**Fig. 4B**; **Fig. 4D, blue shade**). δ decreases at AP #2 and remains low throughout the train, explaining LFD. ρ remains close to its resting value of 0.9, so that the RS gate is closed before each AP of the train (**Fig. 4B, bottom**; note that this figure shows DU parameters just before each AP, and that it does not display the timedependent DU changes during ISIs illustrated in **Fig. 3**). These results show that both at high and at low frequency, the proportions of DU states that are reached after the first ISI are kept without major changes throughout the 8-AP train.

**Figure 4:**
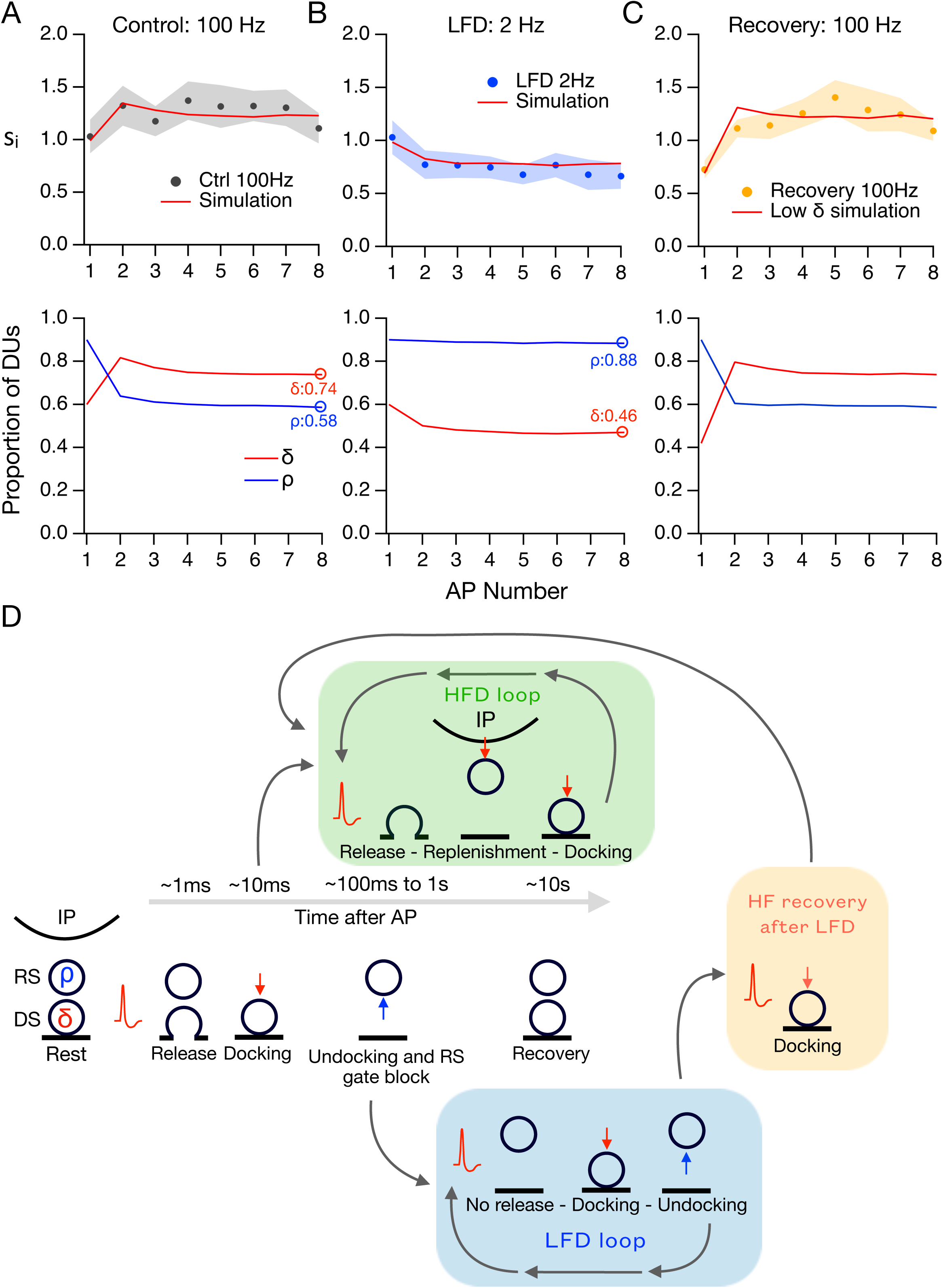
Simulation of SV movements during HFD and LFD. **A, upper panel**: s_i_ curve during control runs with 8 APs at 100 Hz (black dots: data from Fig. 2; red curve: simulated results; model parameters in **Table 1**). **A, lower panel**: simulated values of δ (red) and ρ (blue) observed just before AP stimulations, as a function of AP #. **B**: Same as **A**, but during 2 Hz 8-AP stimulations (data from Fig. 2). Note the behavior of δ and ρ compared to **A**, here ρ remains stable throughout the train, while δ undergoes a sharp initial decrease. At the end of the train there is a low δ value and a high ρ value for 2 Hz trains, while at the end of a 100 Hz 8-AP train (**A**) there is a relatively high δ and a low ρ. **C**: Same as in **A**, but during recovery (100 Hz; data from Fig. 2). The initial δ and ρ values for the recovery train simulation are the end-of-train values from the 2 Hz simulation. **D:** Proposed SV movements during high-frequency depression (HFD) and LFD. **Central sequence:** Proposed timeline of SV movements inside a DU following an AP stimulation. **HFD loop**: Proposed sequence of SV movements during HFD. The DU exits the main timeline at 10 ms after the AP to enter a loop with repetitive high frequency AP stimulation, which results in high rates of exocytosis and RRP replenishment, as the RS gate is open (green shade). Eventually, SV depletion leads to depression. **LFD loop**: proposed sequence of SV movements during LFD. Here the DU exits the main timeline as the RS gate is closed, and it undergoes an idle docking-undocking cycle at each ISI (blue shade). **HF recovery after LFD** proposed sequence of SV movements during high frequency recovery train after LFD (yellow shade). The SVs that were in the RS can dock if high frequency is applied at any point during an LFD train; if further stimuli are applied, the DU is transferred to the HFD loop (green shade) after the 2nd AP.

The simulation of the recovery train used the same transition parameters as the control train. The initial ρ and δ were obtained from simulation values at the end of LFD (**Fig. 4B, bottom**). As can be seen by comparing **Fig. 4C, bottom** with **Fig. 4A, bottom**, DUs behave identically to control from AP #2 to AP #8, in agreement with the data. Thus, our model predicts that a synapse where δ has decreased following LFD can return to its basal state within a single ISI of 10 ms when challenged with a high frequency stimulation (**Fig. 4D, yellow shade**). If a high frequency train is continued the synapse enters the HFD loop (**Fig. 4D, green shade**).

### LFD does not require previous SV release events

The results and simulations of Figs. 2-4 suggest that LFD reflects a drop in δ, but they leave the question open as to whether the δ decrease is due to exocytosis, to SV return to RS (undocking), or to both. To better appreciate the share of each of these two mechanisms in LFD, we re-examined LFD experiments depending on the value of s_1_. If s_1_ is nil, the first theory predicts no synaptic depression in response to the 2^nd^ AP, because there was no prior loss of SVs by exocytosis. Conversely, a failure of release after AP #1 should not prevent synaptic depression according to the second theory. We set aside EPSC sequences during LFD where the 1^st^ AP resulted in release failure (s_1_ = 0; labelled with a star in **Fig. 5A**). We pooled together results obtained with ISI values of 500, 800 and 1600 ms, as the PPR did not appear to depend on the ISI value in this range (PPR = 0.74 ± 0.05, 0.88 ± 0.04, and 0.75 ± 0.04 for ISIs of 500 ms, 800 ms and 1600 ms respectively). We calculated separately mean s_2_ values for the failure traces (noted <s_2_>Is_1_=0) as well as overall mean s_2_ values (noted <s_2,all_>). We obtained two separate LFD values (LFDfail and LFDall) by dividing <S2>Is1=0 and <s2,all> with the overall s1 mean, <s1,all>. With this normalization procedure, the normalization factor was the same for successes and for failures. The two LFD values were significantly < 1 (p = 0.004 for LFDfail and p < 0.0001 for LFDall, one-tailed, one-sample Wilcoxon rank test). Strikingly, we found that the two LFD values were similar (LFD_fail_ = 0.84 ± 0.05 vs LFD_all_ = 0.79 ± 0.03; p = 0.24, unpaired onetailed Wilcoxon rank test; **Fig. 5B, left**), indicating that the value of the LFD does not depend on whether the 1^st^ AP leads to a failure or to a response.

**Figure 5:**
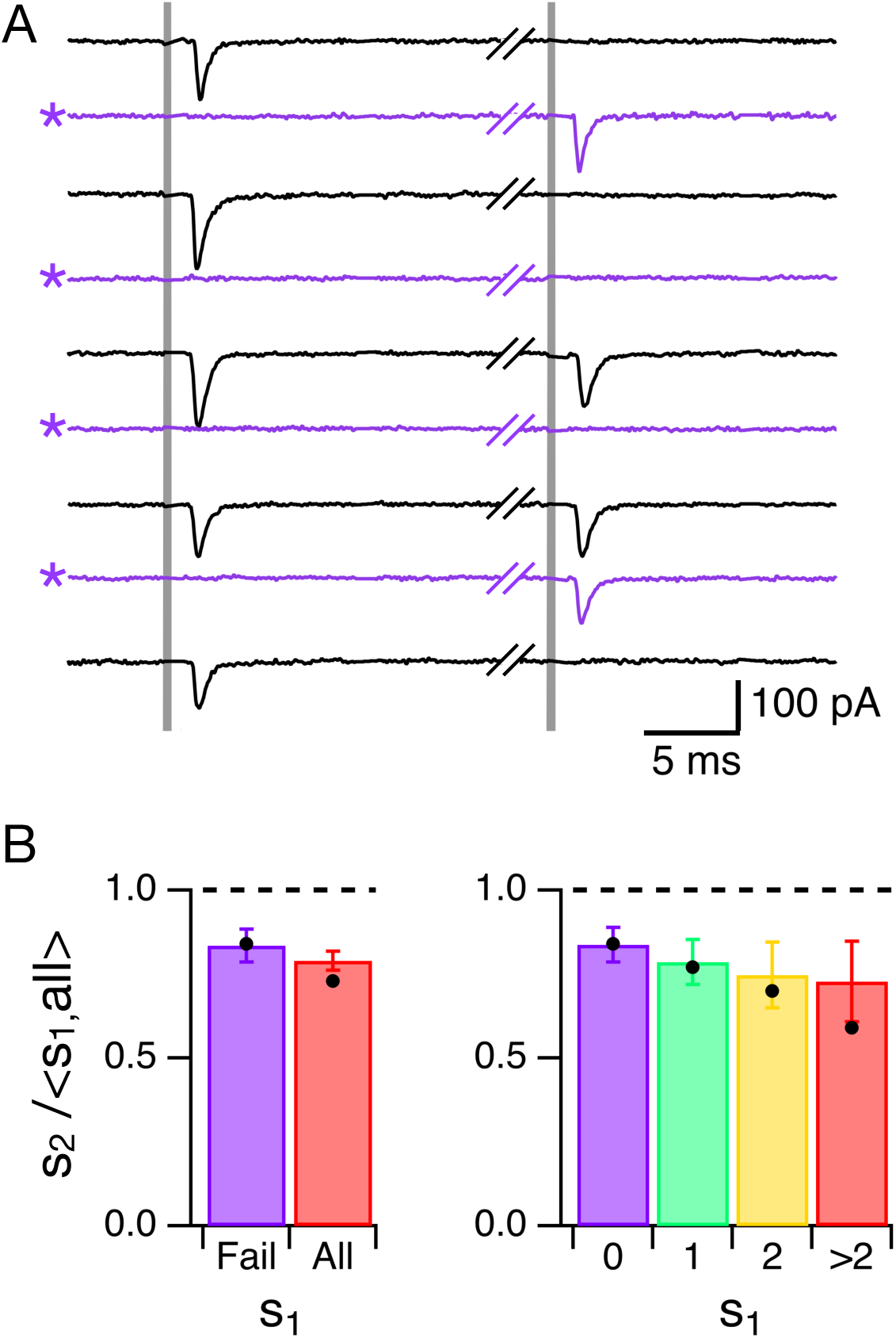
RRP depletion does not cause LFD. **A**: Exemplar traces from an LFD experiment, illustrating responses to 1^st^ and 2^nd^ AP (AP times indicated by vertical grey lines) during low-frequency trains. Black traces show EPSCs including a success in response to the 1^st^ AP, while purple traces (marked with a star) show EPSCs when there was a failure in response to the 1^st^ AP. **B, left**: Group data showing a similar extent of LFD (mean ± sem of LFD across cells; n = 7 cells for 500 ms; 8 cells for 800 and 1600 ms) when there was a failure in response to the 1^st^ AP (purple bar; m. ± sem) compared to the corresponding data taken from all traces (red bar). The LFD value was obtained by calculating the ratio of the mean numbers of released SVs for the second AP over the average release for the first AP for all trials (failures and successes). **B, right**: Same analysis, except that now LFD values are calculated separately for each s_1_ value. Dots indicate predictions from the model of Fig. 3.

Next, we expanded the above analysis by sorting traces displaying successful EPSCs as a function of s_1_. Experimental LFD values were < 1 in all cases: 0.79 ± 0.07 for s_1_ = 1; 0.75 ± 0.10 for s_1_ = 2; 0.73 ± 0.12 for s_1_ > 2; respective one-tailed p values for LFD < 1: 0.002, 0.01 and 0.03 (**Fig. 5B, right**). We finally asked whether these results were compatible with our undocking model. Simulation results using the model of **Fig. 3** gave: LFD_fail_ = 0.84; LFD_all_ = 0.73; LFD_s1 = 1_ = 0.77; LFD_s1 = 2_ = 0.70; LFD_s1 > 2_ = 0.59. These values are close to experimental data (**Fig. 5B**, black dots vs. bars). This agreement is remarkable since the model predicted LFD values without any parameter adjustment. Notably, the finding of identical experimental and simulated LFD_fail_ values (0.84 in both cases), is compatible with the undocking mechanism of **Fig. 3** for LFD.

When comparing the evolution of DUs after SV release vs. after failure in the RS/DS model, it appears that the difference in DS occupancy resulting from release in the first case is compensated in a time frame on the order of 10 ms due to the high value of the RS -> DS transition rate. Because of this short recovery time, DUs that have released display very similar δ (before the 2nd AP) to those that have failed to release after the 1st AP, leading to almost identical s2 values in the two cases.

In conclusion, the above results indicate that LFD occurs independently of the amount of SV release following AP #1. They support the hypothesis that LFD reflects a shift of SV occupancy favoring RSs over DSs (undocking), rather than RRP depletion caused by exocytosis.

### Long AP trains at low frequency reveal two phases of LFD occurring on widely different time scales

Earlier studies showed a slow onset LFD during long trains at low frequency (Abrahamsson et al., 2007; Doussau et al., 2010; Rudolph et al., 2011). We therefore searched for an additional slow component LFD at PF-MLI synapses. In each experiment, a series of control trains (8-APs @ 100 Hz) was presented first. Next, a long low frequency train was applied (200 APs @ 2 Hz in **Fig. 6A**), followed by a recovery train (50 or 100 APs @ 100 Hz; **Fig. 6B**; protocol in **Fig. 6A, inset**). Numbers of released SVs per AP (s_i_) were normalized with respect to the s_1_ value obtained for the control trains. During the long train, a plot of the normalized s_i_ (indicated as s_i_/ctrl s_1_) vs AP # displayed an initial decrease followed by a gradual depression (**Fig. 6A**). We call this initial drop and the subsequent linear decrease the first and second phase of LFD respectively. A linear fit to these data (red line in **Fig. 6A**) indicated a first phase depression (within the first 10 APs) to 75 ± 2 % of the control, consistent with the previous results obtained with paired stimulation and short train protocols (**Figs. 1 and 2**). The second phase of the depression exhibited a slope of -0.15 ± 0.02 % per AP, such that the value reached for the linear fit at the end of the 200-AP train was 45 % of the control. A similar LFD was seen at 1 Hz with a first phase drop to 51 ± 2 % of control and a slope of -0.05 ± 0.02 % reaching 40 % of the control at the end of the 200-AP train (n = 17 trials from 6 cells).

**Figure 6:**
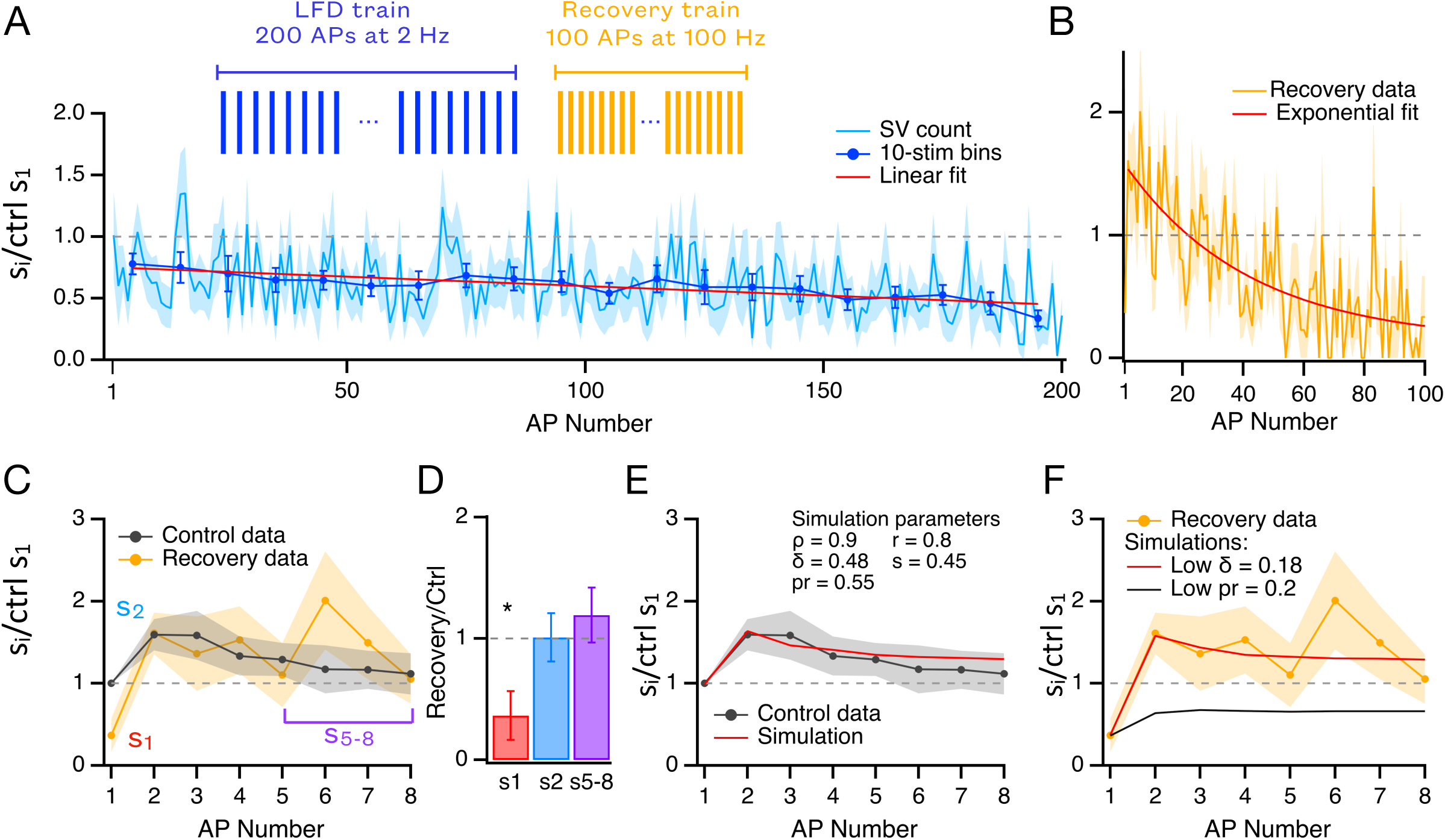
Prolonged low frequency trains produce a gradual decrease in docking site occupancy. **A**: A long AP train at low frequency (200 APs @ 2 Hz) was followed by a long recovery train (50-100 APs @ 100 Hz; experimental protocol in **insert**). Normalized plot of s_i_ during long low frequency train, showing an initial rapid depression (as in Fig. 2, notice the reduction in release for the first binned data point) followed by a gradual depression during the entire train duration (light blue: mean ± sem of individual trials; dark blue dots and associated error bars: binned data for 10 consecutive APs; red: linear fit to the data; n = 16 trials, 8 cells). **B**: Time course of synaptic depression during the recovery train, with exponential fit (red; time constant: 41.2 ISIs, or 412 ms). n = 6 trials, 3 cells for 50 APs, 6 trials, 4 cells for 100 APs. **C**: A series of control trains (8-APs @ 100 Hz, separated by 10 s-long inter-train intervals) were recorded before the long AP train to establish the characteristics of the synapse. Normalized s_i_ plots for this control and the first 8-APs of the recovery 100 Hz train are compared here. s_1_, s_2_ and s_5-8_ are indicated in the plot. **D**: The s_1_ ratio (red) between recovery and control trains is < 1, indicating a decrease in δ at the end of the low frequency train. By contrast, neither s_2_ (blue) nor s_5-8_ (purple) are changed, suggesting no change in ρ or in IP size. **E**: Simulation of control trains shown in **C** (black dots: average normalized data; red curve: simulation). **F**: Simulation of recovery trains shown in **C** (yellow dots: average normalized data). Decreasing the release probability from p_r_ = 0.55 to p_r_ = 0.2 without changing the other simulation parameters fails to provide a satisfactory fit of recovery data (black curve), while decreasing the DS occupancy from δ = 0.48 to δ = 0.18 without changing the other simulation parameters provides a satisfactory fit of the recovery data (red curve).

We next investigated δ, ρ and IP pool size at the end of long low frequency trains with the previous s_1_/ s_2_/ s_5_-s_8_ approach. As expected, the first response of the recovery train was severely depressed compared to s_1_ of control trains (recovery s_1_ = 37.6 ± 20.2 % of control s_1_), but s_i_ plots during control and recovery trains were indistinguishable for i values of 2 or more (s_2_ ratio: 1.01 ± 0.20; s_5-8_ ratio: 1.19 ± 0.23; **Fig. 6C-D**), suggesting again a drop in δ without changes in either ρ or IP size.

The s_1_/ s_2_/ s_5_-s_8_ analysis assumes a constant p_r_. To test this hypothesis, we determined the parameters of the RS/DS model by fitting responses to control 8-AP trains (**Fig. 6E**). Next, responses to the first 8 APs of the recovery train were predicted by keeping all parameters the same as control except for either δ or p_r_. In two different simulations, δ and p_r_ were determined to fit the s_1_ point of the recovery curve (**Fig. 6F**, red and black curves respectively). While the δ decrease prediction closely matches the data (**Fig. 6F**, yellow curve), the p_r_ decrease prediction fails to simulate the recovery train. Altogether, these results suggest that the slow phase of LFD illustrated in **Fig. 6A**, like the rapid phase shown in **Fig. 2C**, reflects a δ decrease without significant changes in p_r_, in ρ or in IP size.

In these experiments the delay between the LFD train and the recovery train was variable. A plot of the value of s_1_ for the recovery plot as a function of the time interval since the end of the LFD train is shown in **Supplementary Figure 4**. This plot shows that recovery s_1_ remains depressed after rest periods of up to 1 min after the end of the LFD train. These results suggest that, while SV re-equilibration within the RRP occurs very rapidly under elevated calcium concentration, as presumably occurs during the recovery trains, they occur very slowly under resting calcium concentration.

Finally, a full recovery train (50 or 100 APs @ 100 Hz) elicited a deep synaptic depression (high frequency depression, or HFD; **Fig. 6B**), similar in extent and in time course to the depression observed for stand-alone trains at high frequency. An exponential fit to these data (red curve in **Fig. 6B**) displayed a time constant of 412 ms (41.2x the ISI) and an asymptotic value of 13% of the control s_1_ value. This type of depression, unlike LFD, was previously suggested to involve a decrease in ρ and in IP size in addition to δ (Tran et al., 2022).

### LFD is not due to stimulation failures

One tacit assumption of our analysis so far is that LFD reflects a decreased synaptic response to presynaptic stimulations. We next asked whether LFD could instead originate in erratic stimulation failures. This possibility was plausible since the failure rate increased markedly during the second phase of LFD (**Supplementary Figure 5A**). To test it, we compared the changes in the distributions of released SV numbers predicted from two LFD models. In the first model, during LFD, each DU releases less SVs when stimulated, presumably due to a reduction in δ. In the second model, a fraction of the stimulations fails to elicit a presynaptic AP during LFD, but the probability of release for each stimulated synapse remains constant (**Materials and Methods**). Both models were constrained to reproduce the observed mean number of failures during LFD. Observed distributions of released SV numbers during LFD were compatible with the decreased δ model but not with the stimulation failure model (**Supplementary Figure 5**). We conclude that stimulation failures cannot account for LFD, consistent with the above interpretation that LFD reflects a drop in δ, as well as with previous results in other preparations (Saviane et al., 2002; Silverman-Gavrila et al., 2005)

### LFD is not blocked by postsynaptic BAPTA

At synapses between PFs and Purkinje cells, low frequency stimulation leads to a gradual synaptic depression that has been attributed to a transsynaptic mechanism (Casado et al., 2000). Since the simple synapse method is based on SV counts rather than EPSC amplitude, there is no doubt that the expression of LFD at PF-MLI synapses is presynaptic. Nevertheless, as AMPAR activation elevates the postsynaptic Ca^2+^ concentration (SolerLlavina and Sabatini, 2006), we included BAPTA (10 mM) in the MLI recording solution to test the possibility of a retrograde LFD mechanism that would be driven by postsynaptic Ca^2+^. We applied a long 2 Hz train (100 APs @ 2 Hz) followed by a short 8-AP recovery train (experimental protocol in **Supplementary Figure 6**, insert). The result is very similar to **Fig. 6** data (compare **Supplementary Figure 6** to **Fig. 6A** and **C**) with a first phase drop to 65 ± 3 % of the control and a second phase slope of -0.2 ± 0.05 % per AP. In this case recovery trains were applied immediately following the LFD train (whereas waiting times of 10 s of s were included in the experiments of **Fig. 6**), and the recovery is identical to that observed in **Fig. 6C**, suggesting likewise a reduction in δ with no associated change in ρ or IP size.

The similarity of results with and without a postsynaptic calcium buffer is consistent with a purely presynaptic mechanism for LFD.

### LFD is not due to decreased Ca^2+^ transients

Our recovery results and simulations suggest that LFD is due to a decrease of δ, not of p_r_ (**Fig. 6F**). In line with this notion, earlier work reported no change in Ca^2+^ transients at PFMLI synapses for paired stimulations at short ISIs (Brenowitz and Regehr, 2007; Malagon et al., 2020; Miki et al., 2016). Nevertheless, it seemed possible that a different situation would prevail with multiple stimulations and/or at longer ISIs.

To explore potential changes of presynaptic Ca^2+^ signaling during AP trains, we preloaded individual granule cells with the calcium dye OGB6F (1-2 mM in pipette solution; duration of preloading: 90 s; (Rebola et al., 2019)). After allowing diffusion of the calcium dye in the axon, we imaged single PF varicosities using two-photon laser microscopy (**Fig. 7A**, and **Supplementary Information**), and we measured Ca^2+^-dependent changes in fluorescence during 50-AP train stimulations at 1 Hz, or 200-AP train stimulations at 2 Hz, using stimulation conditions similar to those of the simple synapse experiments of **Fig. 6**. To avoid photodamage and photobleaching, imaging was interrupted after the 5th AP in a train, and it was resumed for the last 5 APs of the train. Traces were averaged from 3-7 trials in each varicosity to improve the signal-to-noise ratio (**Fig. 7A2**). Averaged traces across varicosities revealed little changes in Ca^2+^-dependent fluorescence transients during trains either at 1 Hz (**Fig. 7B**) or at 2 Hz (**Fig. 7C**). DF/F_0_ values for 10 varicosities from 8 granule cells at 1 Hz were: 19.1 ± 3.9 % for the 1st AP; 20.4 ± 2.8 % for the 2nd AP; 19.3 ± 2.8 % for the last AP; for 9 varicosities from 6 granule cells at 2 Hz: 19.4 ± 2.9 % for the 1st AP; 17.9 ± 1.6 % for the 2nd AP; 16.4 ± 1.9 % for the last AP; p > 0.05 in 1 vs. 2 and 1 vs. last, paired comparisons at both frequencies, onetailed Wilcoxon Rank test (**Fig. 7D**). These data indicate that Ca^2+^-dependent fluorescence transients were the same, within the limits of sensitivity of our measurements, in response to the first, the second, or the last AP in a train. In the same data set, we found no difference between basal fluorescence levels at the end vs. the beginning of the trains at 1 Hz (respectively, 107 ± 12 a. u. vs. 103 ± 10 a. u.; p > 0.05, one-tailed Wilcoxon paired rank test; black markers in **Fig. 7E1**), but we found an augmentation of basal fluorescence at the end of the trains compared to the pre-stimulation value at 2 Hz (118 ± 10 a. u. vs. 111 ± 9 a. u.; p = 0.026, one-tailed Wilcoxon paired rank test; blue markers in **Fig. 7E1**). At 2 Hz, after the end of the train, the fluorescence trace relaxed to near its pre-stimulation value with a time constant of 750 ms, confirming that the basal Ca^2+^ concentration had increased during the train (**Fig. 7E2**).

**Figure 7:**
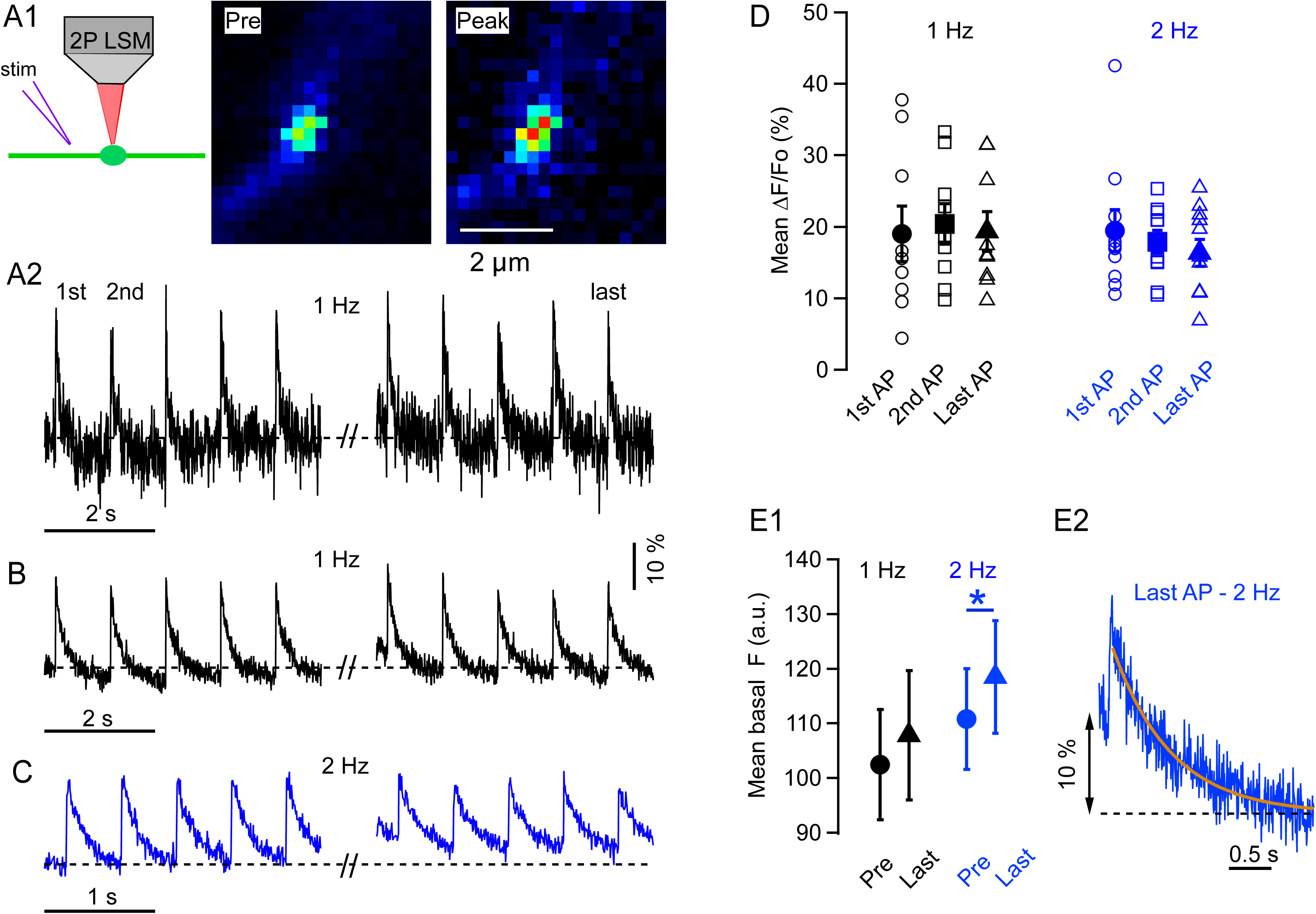
Presynaptic calcium transients in single PF varicosities. **A1, left**: Diagram of the experimental recording. **A1, center:** Image of an OGB-6F loaded PF varicosity before stimulation. **A1, right**: Same, at the peak of the response to a single presynaptic AP. **A2**: Average Ca^2+^-dependent fluorescence transients (DF/F_0_ traces from n = 5 trials: see Methods) registered in response to the 5 first APs (left trace) and to the 5 last APs (right trace) in response to a train of 50 APs at 1 Hz, from the same varicosity as in A1. Imaging was interrupted between the two traces to avoid photodamage and photobleaching. **B**: Average traces from a group of 10 varicosities from 8 cells at 1 Hz. **C**: Average traces from another group of 9 varicosities from 6 cells, in response to the first and last 5 APs from a train of 200 APs at 2 Hz. **D**: Mean Ca^2+^-dependent fluorescence transients (closed symbols: means across varicosities, with attached bars representing ± the sem; open symbols: values for individual varicosities) are the same for the 1st, 2nd, and last AP in a train (left: 1 Hz stimulation; black; right: 2 Hz stimulation; blue). **E1-E2**: Evolution of the basal fluorescence level during and after a long AP train. **E1**: Initial and final basal fluorescence levels are not statistically different for long trains at 1 Hz (left, black), while there is a significant increase at the end of trains at 2 Hz compared to its initial value (right, blue). **E2**: Return of the DF/F_0_ fluorescence trace to its initial baseline (mean from 5 varicosities) after the end of long 2 Hz trains indicates a shift of the basal fluorescence level by 11 % near the end of the train (yellow curve: exponential fit to the decay, with time constant 750 ms).

In conclusion, we find no evidence of decreased Ca^2+^ transients during trains, either at 1 Hz or at 2 Hz. This is in line with our earlier conclusion that p_r_ changes do not contribute strongly to LFD. We find a gradual increase of pre-stimulation Ca^2+^ concentration with 200 APs at 2 Hz, but not with 50 APs at 1 Hz. This effect may contribute to the second phase of LFD at 2 Hz.

### Doublet stimulations inhibit LFD

Our model predicts a high ρ value during LFD that inhibits DU replenishment by preventing the transition from the IP to the RS (RS gate closed). We next tested the prediction that during a low frequency train, ρ remained elevated while δ decreased (**Fig. 4B**), by giving two successive stimuli with a 10 ms interval instead of single stimuli (doublet stimulations; **Fig. 8A-B**). According to the s_1_/ s_2_/ s_5-8_ analysis, while the time course of the first response, s_1_, should give the evolution of δ, the time course of the second response, s_2_, should give the evolution of ρ (Tran et al., 2022). With the same experiments, we intended to test whether doublet stimulations would open transiently the RS gate and thus interfere with the time course of LFD, as suggested by earlier work at the PF-Purkinje cell synapse (Doussau et al., 2017).

**Figure 8:**
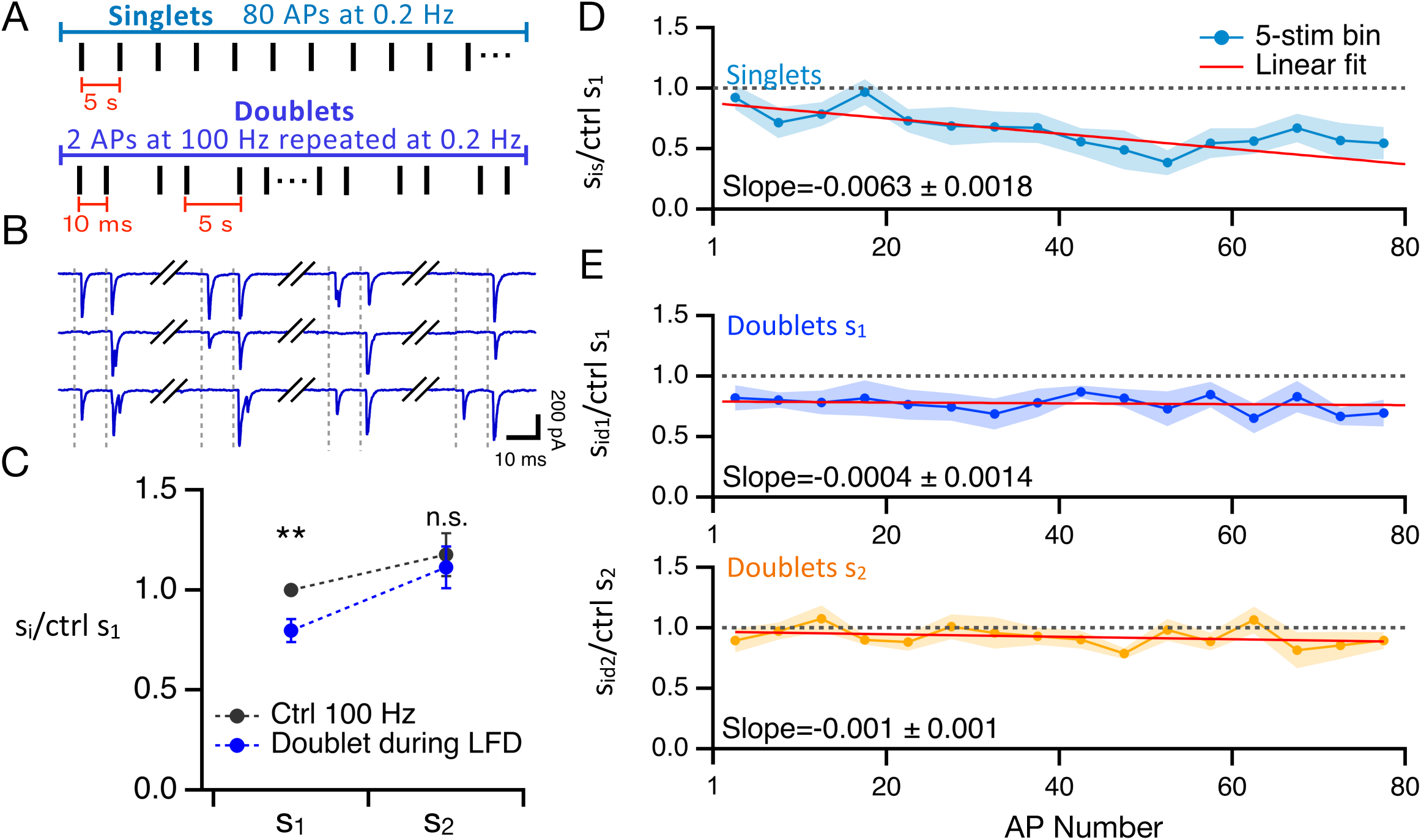
LFD during doublet vs. singlet train stimulations. **A:** Stimulation protocols. Either an 80-AP train of single stimulations (‘singlets’) or a sequence of 80 double stimulations 10 ms apart (‘doublets’) were applied at 0.2 Hz. Having 2 stimuli provides an evaluation of ρ and δ during LFD by comparing release during control and after AP# 1 or AP# 2 during doublets (s_id1_ and s_id2_, respectively). **B:** Example traces of a doublet experiment. **C:** During the first 20 doublet stimulations, the ratio between s_id1_ and ctrl s_1_ is < 1, indicating a decrease in δ; however, s_id2_ is not significantly different to ctrl s_2_ indicating that the occupancy of the replacement sites (ρ) remains the same as control during LFD. **D:** Normalized plot of s_i_ during singlet trains (s_is_) showing a 2-component depression (compare to Fig. 3). The initial component is followed by a gradual steep depression during the entire train duration (mean ± sem of individual trials, normalized with respect to the mean s_1_ value obtained during preliminary control trains; blue dots and associated error bars: binned data for 5 consecutive APs; red, linear fit to the data; 8 trials from 5 cells). **E:** Normalized plot of s_i_ during doublet trains (mean ± sem of individual trials, normalized with respect to the mean value obtained during preliminary control trains; dots and associated error bars: binned data for 5 consecutive APs; red, linear fit to the data; 9 trials from 6 cells). **Top:** SV release for the first AP of each doublet (s_id1_) displays a rapid LFD component but no significant slow component. **Bottom:** SV release for the second AP of each doublet (s_id2_), showing no significant deviation from the control value.

We performed these experiments with a longer than usual ISI (5 s) to provide ample time for SVs to equilibrate after each stimulus, either singlet or doublet. Control single AP train stimulation experiments at 0.2 Hz revealed LFD (0.75 ± 0.11; paired one-tailed t-test, p = 0.037, n = 8; exponential fit of first 12 normalized si points had a time constant of 3.6 ISIs, or 18 s). We then examined the extent of LFD in response to the first 20 doublet stimulations at low frequency. We found that s_1_ values were reduced compared to control (normalized s_1_ value during 20 first trials at 0.2 Hz: 0.80 ± 0.10; p = 0.008, two-tailed, one-sample t-test; **Fig. 8C**). These results indicate that doublet stimulations at 0.2 Hz produced significant LFD when considering the response to the 1^st^ stimulus. By contrast, s_2_ values were not affected (s_2_ values normalized to control s_1_: 1.18 ± 0.11 for control 100 Hz trains; 1.11 ± 0.10 during 20 first doublet trials; p > 0.05, two-tailed, one-sample t-test; **Fig. 8C**). These results are consistent with our previous conclusion that LFD involves a decrease in δ without significant changes in ρ.

Next, we compared the plots of normalized released SV numbers as a function of AP number (i) for singlet stimulations (s_is_; light blue), for the 1^st^ AP of doublet stimulations (s_id1_; dark blue), and for the 2^nd^ AP of doublet stimulations (s_id2_; orange; **Fig. 8 D-E**). In each case, the curves could be approximated to linear decay curves as a function of i (y = α – βi), but the line parameters differed for the 3 curves. Values of α were similar for s_id1_ and for s_is_ (0.79 ± 0.07 and 0.88 ± 0.11 respectively; p = 0.53, two-tailed, two-sample t-test). This indicates that the first phase of LFD occurs with a similar amplitude for singlets and for doublets. On the other hand, values of β were markedly smaller for s_id1_ than for s_is_ (β_d1_ = 0.0004 ± 0.0014 vs. β_s_ = 0.0063 ± 0.0018; p = 0.011, two-tailed, two-sample t-test; **Fig. 8D** and **Fig. 8E, top**). These results suggest that the second phase of LFD is less pronounced for doublet stimulations than for singlet stimulations, with a slope ratio near 15-fold. The results for d_2_ can be interpreted as a return of ρ values close to control at the end of each 5 s ISI throughout the train (orange curve in **Fig. 8E, bottom**; α_d2_ = 0.97 ± 0.04; β_d2_ = 0.001 ± 0.001), which suggests that the RS gate was closed at the end of each ISI after transitorily opening during the preceding doublet.

Overall, the responses to the first stimulations in doublets display less LFD than singlets, and the responses to the second stimulations in doublets display no LFD at all. The results are consistent with the notion that LFD involves a closing of the RS gate, which is transiently alleviated by doublet stimulations.

## Discussion

In the present work, we use the SV counting method at simple PF-MLI synapses (Malagon et al., 2016) to investigate cellular mechanisms underlying LFD. Our results reveal similarities, but also significant differences, between these mechanisms and those previously described for moderate or high frequency synaptic depression (Lin et al., 2022; Tran et al., 2022). Based on our results and on our simulations using the RS/DS model, we propose that LFD reflects a decrease in δ following SV undocking. We also explain how the mechanisms of synaptic depression may vary as a function of stimulation frequency.

### PPR as a function of ISI

A traditional view of synaptic physiology has been that synapses may be either facilitating or depressing. This view is clearly an oversimplification, however, as many synapses display a combination of facilitating and depressing properties (Dittman et al., 2000). Here we show that PF-MLI synapses exhibit PPR values > 1 for ISI values comprised between 10 and 200 ms, and PPR values < 1 for ISI values comprised between ∼500 ms and ∼1500 ms (**Fig. 1**). Importantly, the shift from facilitation to depression is apparent not only in group results (**Fig. 1D**), but also in individual experiments (**Fig. 1C**). A similar mix of facilitation at short ISI and depression at long ISI has been described in several central synapses (Chiu and Carter, 2024; Doussau et al., 2017). Following our analysis of simple synapse recordings, and in view of zapand-freeze results in hippocampal cultures, we propose that facilitation and LFD are linked together in a docking-undocking sequence following AP stimulation (Kusick et al., 2020).

### SV movements during HFD and LFD

Based on the present study and on earlier work (Kusick et al., 2020; Miki et al., 2016; Tran et al., 2022), we propose a global model for HFD and LFD as outlined in **Fig. 4D**. If an AP train is presented with short ISIs ( ∼ 10 ms), a rapid sequence of release/docking/replenishment occurs (open RS gate). The arrival of a new AP induces forward movement of SVs (from DS to exocytosis, from RS to DS, and from IP to RS). If a docked SV fails to be released, it does not undock, because a new SV rapidly occupies the corresponding RS. These events lead to facilitation at first. IP depletion eventually reduces SV influx into the RRP, such that the IP size becomes rate limiting at the end of the train. This accounts for HFD (HFD loop in **Fig. 4D**, green shade). If an AP train is presented with longer ISIs (0.5 s to 5 s), by contrast, release and replenishment are reduced since many DUs have their DS empty and their RS occupied at the time of the AP (closed RS gate). During the closed RS gate period, ρ is ∼0.9, so that the entry rate of SVs from the IP into the RRP, which is proportional to (1-ρ), is reduced 10-fold by RS occupation. By comparison, ρ is ∼0.5 when the RS gate is open. The larger ρ value results in a net 5-fold slower entry rate when the RS gate is closed compared to when it is open. With longer ISIs, SVs move from the RS to the DS after each AP, but they have the time to move back to the RS before the next AP. Therefore, DUs tend to undergo an idle docking/undocking loop without release, leading to LFD (LFD loop in **Fig. 4D**, blue shade).

AP induced docking and undocking happen in overlapping time frames. In **Fig. 1** the highest amount of docking is seen at ∼10 ms ISI (it is also present at 5 ms: (Miki et al., 2016; Silva et al., 2024)). Undocking on the other hand is maximal more than 1 s after the AP. For ISIs in between there is a mix of docking and undocking. According to this model, the PPR recovery observed at ISIs of ∼200 ms is only apparent because in this period undocking is still ongoing.

When stimulation is switched to high frequency after LFD, SV release recovers within 10 ms to the same level as during high frequency control trains (**Fig. 2D**; **Fig. 6C**). According to the RS/DS model, this happens because the SVs that are in the RS dock with each AP and are still in the DS when a second AP is presented rapidly (HF recovery in **Fig. 4D**, yellow shade). This would explain why the s_2_ of a recovery train is equivalent to the control value. If high frequency stimulation is continued the synapse will behave identically to a naïve synapse for the remainder of the train (**Fig. 2D**; **Fig. 6C**) and it will eventually undergo HFD (**Fig. 6B**; HFD loop in **Fig. 4D**, green shade).

It has long been recognized that synaptic activity enhances RRP replenishment (Neher and Sakaba, 2008). While a Ca^2+^-dependent rate of SV entry into the RRP has been invoked as underlying mechanism, other explanations cannot be ruled out (Miki et al., 2020; Ritzau-Jost et al., 2018; Ritzau-Jost et al., 2014). Transient opening of the RS gate, as suggested here, offers a novel mechanism explaining activity-dependent RRP replenishment.

### LFD may involve a specific δ decrease rather than across-the-board RRP decrease

Our conclusion that LFD results from a δ decrease is largely based in our analysis of high frequency recovery trains. While this approach is model-dependent, as it assumes the RS/DS model of **Fig. 2A**, our conclusion is supported by two additional lines of evidence. First, we find that LFD occurs even after failures, and that the extent of LFD does not depend significantly on s_1_ (**Fig. 5**). These findings argue against the standard RRP depletion mechanism. By contrast, the docking-undocking sequence of **Fig. 4D** (main sequence and LFD loop) accounts for our results as it predicts that the DU state at the end of a long ISI depends little on the response to the previous stimulation (simulation in **Fig. 5B**). Secondly, we find that LFD is alleviated when using doublet rather than singlet stimulations (**Fig. 8**). Again, this result is inconsistent with an across-the-board RRP depletion model, but it can be readily explained by the RS/DS model. During LFD with singlet stimulations, the RS/DS model predicts a gradual δ decrease accompanied by a high ρ (**Fig. 3**). Doublet stimulations change this pattern in two ways. Firstly, they transiently lift the RS block that prevents RRP refilling with singlet stimulations, leading to enhanced RRP replenishment. The second AP of the doublet arrives in the period where SVs are still present in the DS, releasing a fraction of them, and promoting SV movement from the RS to the DS. This means that SVs coming from the IP are allowed, in turn, to bind the now-free RS. Secondly, doublet stimulations prevent SV undocking. This happens as the incoming SV at the RS locks the SV that is bound to the DS in its docked position. The combination of these two effects leads to an inhibition of the slow phase of LFD during doublet stimulations. Of note, this complex RS/DS interaction is only possible because the RS/DS model allows simultaneous occupation of both RS and DS by distinct SVs (Silva et al., 2024).

Altogether, the present LFD model resembles previous studies assuming two different SV states inside the RRP, and a decrease in the percentage of primed/docked SVs during synaptic depression (Doussau et al., 2017; Tanaka et al., 2021; Tran et al., 2022; Lin et al., 2025). It differs from these earlier studies in proposing a role of RS in gating SV movements during LFD.

While our model assumes that the δ decrease associated with LFD is primarily due to the return from DS to RS (undocking), we cannot exclude alternative mechanisms. SVs could leave the DS following exocytosis, or they could unbind and diffuse away in the cytosol. A previous study suggested that delayed release is insufficient to account to the δ decrease accompanying the end of facilitation (Miki et al., 2018), making the first option unlikely, but it remains possible that during LFD, part of the δ decrease is due to SV escape into the cytosol independently of the DS->RS transition.

### Alternative presynaptic depression mechanisms

Simple synapse recording and deconvolution analysis give us a direct estimate of the number of SVs released, bypassing any possible postsynaptic effects. Expression of LFD as described in the present work is therefore presynaptic. Furthermore, our BAPTA experiments (**Supplementary Fig. 6**) argue against a participation of the postsynaptic compartment to LFD induction, so that both induction and expression of LFD are likely presynaptic. We consider here other presynaptic mechanisms than undocking that have been proposed to underlie LFD. A p_r_ reduction has been suggested (Wölfel et al., 2007; Wu and Borst, 1999), either resulting from a direct reduction in Ca^2+^ current (Xu and Wu, 2005) or from the buffering effect of calretinin (Bolshakov et al., 2019). The rapid reversal obtained during recovery trains in the present work is not consistent with either of these hypotheses, as both Ca^2+^ channel inhibition and a buffering effect would require hundreds of ms to seconds to recover. Furthermore, the results and simulations shown in **Fig. 6F** suggest a change of δ but not of p_r_ during LFD. Finally, our Ca^2+^ measurements in presynaptic varicosities do not suggest a difference in Ca^2+^ entry following repetitive low-frequency stimulation although there is a change in basal calcium during long trains that could be linked to the second, slow phase of LFD (**Fig. 7**). Of note, Ca^2+^ measurements have also been performed in relation to LFD in neocortical L4-L2/3 synapses with synaptic depression and no change in Ca^2+^ signal for paired stimuli was reported at ISIs of 500 ms (Chiu and Carter, 2024). In *Aplysia*, silencing of docking sites was suggested as the reason behind synaptic depression (Doussau et al., 2010), however a decrease in N would not predict the rapid return of high frequency release when recovery trains are applied after LFD.

In conclusion, it is unlikely that a p_r_ or N reduction underlies LFD. Overall, a δ reduction best explains LFD and its fast recovery.

### Onset and offset kinetics of LFD

In some studies, LFD has been reported to develop almost immediately, after one or a few stimulations (Chiu and Carter, 2024; Rudolph et al., 2011) whereas in others it develops much more slowly as a function of stimulation number (Abrahamsson et al., 2007; Doussau et al., 2017). In the present work we observe an immediate reduction in synaptic output (**Fig. 2**) followed by a very slow phase which does not reach a steady state after 200 APs at 2Hz (**Fig. 6A**) and 80 APs at 0.2 Hz (**Fig. 8D**). After a short train, in which only the initial phase is observed, and after a long train, where both phases are present, we find a reduction in DS occupancy but no changes in RS or IP occupancy (**Fig. 2E** and **6D**). This leads us to propose that LFD comprises two phases with different onset kinetics.

We find that LFD fails to recover significantly following a rest period of up to 1 minute (**Supplementary Fig. 4**). These results are consistent with earlier studies showing that for sparse stimulations, LFD recovery occurs on a time scale of minutes (Abrahamsson et al., 2007; Doussau et al., 2010; 2017). If, however, a high frequency train is presented, the synapse recovers from LFD almost instantaneously, within one single ISI (range 5-20 ms (Doussau et al., 2010, 2017); present work, **Fig. 2, 6 and 8**). As seen in **Fig. 4D**, the high frequency recovery loop after LFD is identical to the HFD loop: presenting new APs at short ISI does not leave enough time for undocking and immediately brings the synapse back to the HFD mode. Altogether, LFD places the synapse in a depressed state that is stable if stimulation is absent or infrequent, but that is immediately reversed if a high frequency train is presented. In our model, this dichotomy occurs because the SV-RS association is stable at rest, when presynaptic Ca^2+^ concentrations are low (RS gate closed) but is unstable during high frequency AP trains, when presynaptic Ca^2+^ concentrations are elevated (RS gate open). In contrast to LFD recovery, HFD recovers within seconds following the end of a high frequency train, reflecting the gradual replenishment of the IP (Tran et al., 2022).

### Frequency-independent synaptic depression

In the present work, we observe similar extents of LFD at the end of the first phase for stimulation frequencies of 0.2 Hz (0.75 ± 0.11), 1 Hz (0.79 ± 0.08), and 2 Hz (0.69 ± 0.05). These results indicate that at PF-MLI synapses, the first phase of LFD results in a largely frequencyindependent steady state synaptic output in the frequency range 0.2 Hz to 2 Hz. It has been likewise noted in earlier studies that the extent of synaptic depression at steady state is largely independent of stimulation frequency in a certain frequency range, albeit in a higher range than in the present work (10-100 Hz at Purkinje cell to deep cerebellar nuclei synapses (Turecek et al., 2016); 5-50 Hz at the calyx of Held (Lin et al., 2022)). These results have been interpreted either by a compensation between an increased facilitating component and an increased depressing component at higher frequencies (Turecek et al., 2017, 2016), or by a time limitation on calcium-dependent movement of SVs from a primed to pre-primed location, so that this movement is regulated by AP number rather than by time (Lin et al., 2025).

### Molecular mechanisms of docking and undocking

The molecular mechanisms of docking and undocking are under intense scrutiny. Consistent with a role in AP-dependent docking, synaptotagmin 7 (syt7) is associated to facilitation in different preparations (Huson and Regehr, 2020; Jackman et al., 2016; MacDougall et al., 2018). Using zap-and-freeze electron microcopy in hippocampal cultures revealed that syt7 KO showed no rapid docking ∼10 ms after an AP, which was translated into a depressing phenotype when trains of APs are applied (Wu et al., 2024). In neocortical synapses, a comparison of PPR curves in WT and syt7 KO showed no facilitation at short ISIs and a deeper depression at longer ISIs in the KO, with the same recovery time frame (5 s) (Chiu and Carter, 2024), supporting a role of syt7 in AP-dependent docking, but not in undocking or baseline docking dynamics. Recently, intersectin-1 and endophilin A1 were reported to stabilize SVs in the replacement site (RS) (Ogunmowo et al., 2025), indicating a role in docking/undocking and possibly in LFD. Munc-13 is another likely candidate for docking (Kusick et al., 2022), as its deletion reduces the number of docked SVs and changes their relative distance to the plasma membrane (Imig et al., 2014; Siksou et al., 2009). In the calyx of Held, RRP recovery after high frequency facilitation and depression depends on both Munc13 and synaptotagmin 3 (syt3; Weingarten et al., 2022). Modeling suggested that both proteins are required to explain AP-dependent docking.

In conclusion, multiple proteins have been proposed as candidates for AP-dependent docking; in our preparation, Munc-13, syt3 and 7 are likely candidates. By contrast, the molecular mechanisms behind undocking remain largely unknown. It is likely that some of the STP heterogeneity arises from variations in protein expression and regulation amongst synapses, so that the docking/undocking kinetics are not the same. However, LFD is present in multiple preparations, and may be a general mechanism.

### Functional implications

*In vivo* MLI recordings show very infrequent EPSCs in the absence of PF input (< 1 Hz), while sensory stimulation results in short EPSC trains at high frequency (Chadderton et al., 2004; Jörntell and Ekerot, 2003). LFD as described here likely contributes to lower the frequency of background EPSCs, as it inhibits the responses to infrequent presynaptic APs. However, in view of the rapidity of the post-LFD recovery, full EPSC responses are likely restored as soon as a high frequency train of presynaptic APs is presented. This should increase the signal-to-noise ratio in a burst vs. background signaling mode and should improve the sensitivity of the cerebellar circuitry to external inputs. As many brain neurons display low background firing rates (< 1 Hz) *in vivo* (Buzsáki and Mizuseki, 2014; Margrie et al., 2002), a similar signal-to-noise improvement could occur in various brain regions following LFD.

### Materials and Methods Experimental model

C57BL/6 mice (12–16 days old; either sex) were used for preparation of acute brain slices. Animals were purchased from Janvier Laboratories (RRID: MGI: 2670020). They were housed and cared for in accordance with guidelines of Université Paris Cité (approval no. D 75-06-07; animal rearing service of the BioMedTech Facilities, INSERM US36, CNRS UMS2009).

### Slice preparation

200-μm thick sagittal slices were prepared from the cerebellar vermis as follows. Mice were decapitated under anesthesia and the cerebellar vermis was carefully removed. Dissections were performed in ice- cold artificial cerebrospinal fluid (ACSF) which contained the following (in mM): 130 NaCl, 2.5 KCl, 26 NaHCO_3_, 1.3 NaH_2_PO_4_, 10 glucose, 1.5 CaCl_2_, and 1 MgCl_2_. Slices were cut using a vibratome (VT1200S; Leica) and incubated in ACSF at 34°C for at least 60 min before being used for experiments.

### Synaptic electrophysiology

Whole-cell patch-clamp recordings were obtained from MLIs (comprising both basket and stellate cells). The extracellular solution contained (in mM): 130 NaCl, 2.5 KCl, 26 NaHCO_3_, 1.3 NaH_2_PO_4_, 10 glucose, 3 CaCl_2_ (1.5 in the experiments where it is stated), and 1 MgCl_2_ (pH set to 7.4 with 95% O_2_ and 5% CO_2_). The internal recording solution contained (in mM): 144 K-gluconate, 6 KCl, 4.6 MgCl_2_, 1 EGTA (ethylene glycol- bis tetraacetic acid), 0.1 CaCl_2_, 10 HEPES (4- (2- hydroxyethyl)- 1- piperazineethanesulfonic acid), 4 ATP- Na, 0.4 GTP- Na (pH 7.3, 300 mosm/l). Gabazine (3 μM) was included in all experiments to block GABA_A_ receptors. Recording temperature was 32–34°C. To establish a single PF–MLI connection, puffs using a glass pipette filled with the high K^+^ internal solution were applied in the granule cell layer to identify a potential presynaptic granule cell. The same pipette was then used for extracellular electrical stimulation of the granule cell, with minimal stimulation voltage (stimulus voltage was higher than threshold by ∼5 V; range: 10–40 V; 150 μs duration). If the resulting EPSCs, particularly those that occurred after a short stimulation train, were homogeneous, then it was likely that only one presynaptic cell was stimulated. Recordings were only accepted as a simple synapse recording after analysis if the following three criteria were satisfied (Malagon et al., 2016): (1) the EPSC amplitude of the second release event in a pair was smaller than that of the first, reflecting activation of a common set of receptors belonging to one postsynaptic density; (2) all the EPSC amplitudes followed a Gaussian distribution with a coefficient of variation < 0.5; and (3) the number of release events during the baseline recording was stable.

### Quantification and statistical analysis

Ofline analysis of electrophysiological data and statistical analysis was performed using Igor Pro (WaveMetrics). Data are expressed as mean ± standard error of the mean (SEM). Error bars in all graphs indicate SEM. Details of statistical tests are described in the text or figure legends. Student’s t-test was used when the distribution was Gaussian; Wilcoxon Rank test was used otherwise. One-tailed tests were used when the sign of the effect was predicted before performing the experiments; two-tailed tests were used otherwise. Statistical significance was accepted when p < 0.05. The number of trials and cells (some cells have multiple trials) for each experiment is indicated by a lowercase n. Cells were derived from different animals. Each experiment started with control recordings of 8-AP trains @ 100 Hz, with 10 s inter-train intervals. At least 25 trials were performed for this initial control run before proceeding to specific stimulation protocols. For PPR experiments (**Fig. 1**) 4-AP trains were applied with varying ISIs (20, 100, 200, 400, 800, 1600 and 3200 ms) in a scrambled order, with a rest period of 10 s between trains (**Fig. 1A, right**); only synapses with minimum 20 repetitions of this protocol were considered for analysis. For **Fig. 2**, 8-AP trains at alternating frequencies (100 Hz and 2 Hz) were applied. A 100 Hz recovery train was applied immediately after the end of each 2 Hz train (see **Fig. 2B**); a minimum of 20 repetitions of the protocol were obtained for each synapse. Since the value of s_1_ should be independent of train frequency (the synapse cannot predict the frequency at which the train will be given), values of s_1_ were averaged across frequencies in each synapse. Therefore in **Fig. 1D**, the s_1_ SV # used for calculation of the PPR was averaged across 8 frequencies. In **Fig. 2**, the s_1_ value used for normalization is the average of s_1_ values for the control 100 Hz train and for the 2 Hz train at each train cycle. For these two protocols the repetitions of each synapse were averaged together, and synapse results were averaged for group results. For **Fig. 6** a 200-AP train was applied at low-frequency (2 Hz), followed by a 50-AP or 100-AP high-frequency (100 Hz) recovery train (**Fig. 6A**). The time interval between the last AP in the LFD train and the 1st AP in the awakening train varied in the experiments (**Supplementary Fig. 4**). Synapses were considered with minimum one full trial (low frequency + recovery), the maximum number of trials performed in one synapse was 4. Trials were averaged for group results. For **Fig. 8**, 80AP trains were applied at 0.2 Hz with either a single or a double stimulus (**Fig. 8A**), trials were averaged together for group results. In 5 cells singlets and doublets were applied sequentially. For all protocols a new control run was performed when possible at the end of the experiment to verify functional stability of the synapse. For each set of experiments the numbers of repetitions are stated in the figure legend.

### Decomposition of EPSCs

The time of occurrence and the amplitude of individual release events were determined based on deconvolution analysis, as described previously (Malagon et al., 2016). In brief, for each synapse, an mEPSC (miniature excitatory postsynaptic current) template was obtained from delayed release events recorded after the end of AP trains. This template was fitted with a triple- exponential function with five free parameters (rise time, peak amplitude, fast decay time constant, slow decay time constant, and amplitude fraction of slow decay). These five parameters were then used for deconvolution of the template and of individual data traces, producing a narrow spike (called spike template) and sequences of spikes, respectively. Next, each deconvolved trace was fitted with a sum of scaled versions of the spike template, yielding the timing and amplitude of each release event. The amplitude was further corrected for receptor saturation and desensitization, using the exponential relationship between individual amplitudes and the time interval since the preceding release events. Events that were at least 1.7 times larger than the average mEPSC were split into two (Miki et al., 2017). s_1_ was determined as the number of SVs released within 5 ms after the first AP of a train. s_2_ was the number of SVs released within 5 ms after the second AP and s_5-8_ was the cumulative number of SVs released 5ms after each stimulus for AP# 5 to 8. As explained in Tran et al. (2022) (Tran et al., 2022), these 3 parameters are proxies respectively for the occupancy of DS (δ), the occupancy of RS (ρ), and the IP size.

### Calcium measurements

To perform Ca^2+^ imaging of single PF varicosities we pre-loaded granule cells with a calcium indicator. A whole-cell recording was made with a K^+^ gluconate-based solution containing 166 K gluconate, 4.1 MgCl_2_, 9.9 HEPES-K, 0.36 Na-GTP and 3.6 Na-ATP, 1-2 mM of the calcium probe Oregon Green BAPTA 6F. The neurons were held in I-clamp mode with currents of -5 to -30 pA injected to keep the average resting potential close to -90 mV for 1.5 to 2 minutes after which the pipette was removed. Obtaining an outside-out patch with > 5 GW resistance indicated successful pipette removal, and the granule cell was left for approximately 15 minutes to allow for dye diffusion into the axonal compartment. After this waiting time, a 2photon laser scanning microscope (Tan et al., 1999) was used to identify an axonal varicosity. A glass pipette with tip diameter around 1 µm, filled with extracellular saline (in mM: 145 NaCl, 2.5 KCl, 10 K-HEPES, 2 CaCl_2_, 1 MgCl_2_), was introduced into the slice at the depth of the varicosity and placed 10-20 µm from the axon. The stimulation protocol consisted of 1 train of 8 pulses (100 µs duration, 10 to 40 V) delivered at 100 Hz, followed by either 50 stimulations at 1 Hz or 200 at 2 Hz and a new 8 pulse train at 100 Hz applied 30 seconds after the end of the long train. This cycle was performed 3 to 7 times. Fluorescence was acquired at dwell time of 5 ms/frame through raster scans of 5 by 5 µm fields encompassing the axonal varicosity. For each long stimulation train that was applied, we recorded two fluorescence traces. One trace covered the first 5 APs of the train (trace duration: 5 s at 1 Hz stimulation frequency, and 2.5 s at 2 Hz stimulation frequency). We next interrupted laser illumination and scanning (during 40 s at 1 Hz, and during 95 s at 2 Hz) to minimize photobleaching and photodamage. We resumed two-photon imaging near the end of the train, collecting another trace covering the last 5 APs. Time dependent fluorescence signals were analyzed in the pixels encompassing the varicosity and quantified in terms of changes relative to pre-stimulus values (ΔF/F_0_ expressed in %) with software written in the IGOR-Pro programming environment (Wavemetric, Lake Oswego, OR, USA). To avoid possible errors due to tissue movement, focusing was performed twice: just before the first trace, and again before the second trace. To estimate changes in baseline Ca^2+^ concentration, we compared basal fluorescence averages (calculated just before the 1st AP in each trace) for early and late traces across trials for all experiments. This revealed an apparent (non-significant) increase by 4 % of the basal fluorescence at 1 Hz stimulation frequency, and a significant 7 % increase at 2 Hz stimulation frequency (**Fig. 7E1**). In view of this baseline shift (F_Inc_), the second DF/F_0_ trace needed a correction. In this trace, the basal fluorescence used to calculate DF/F_0_ was corrected according to F_0corr_ = F_0app_ – F_Inc_, leading to the relation DF/F_0corr_ = DF/F_0app_ * (1 + F_inc_ /F_0_) where DF/F_0app_ and DF/F_0corr_ are the apparent and corrected values of DF/F_0_ for the late trace, and the ratio F_Inc_/F_0_ is 4 % at 1 Hz and 7 % at 2 Hz. For each train of stimuli, ΔF/F_0_ values were averaged over 5 to 7 repetitions at each varicosity and subsequently data from all varicosities were pooled together to generate the average signal as a function of time from which mean peak values were extracted.

### Model and simulation of SV docking and release

Monte Carlo simulation of two-step SV docking and release (**Fig. 3**) was performed using Igor Pro as described previously (Miki et al., 2016; Silva et al., 2024; Tran et al., 2022). The model is created by assuming a fixed number (N) of docking units (DUs) with 2 binding sites for each DU (RS and DS; **Fig. 2A**), each of which can be empty or occupied by an SV. The probabilities of initial occupancy for DS and RS are δ and ρ, respectively. The probability of SV release after one AP of an occupied DS is p_r_. SV transitions within the DU are characterized by transition probabilities s_f_, s_b_, r_f_, r_b_.

For each series of experiments, control runs (usually 30 trials) were initially performed with stimulations of 8 APs at 100 Hz, interspersed by 10 s-long rest periods. We used previously developed procedures to determine from these data the parameters N, δ (the mean DS occupancy before the 1^st^ AP), ρ (the mean RS occupancy before the 1^st^ AP), as well as the probabilities s_f_, r_f_, and p_r_, all of which were assumed constant throughout the AP train. In this part of the simulation, the back transitions s_b_ and r_b_ were neglected. The simulation yielded the numbers of released SVs (s_i_) as a function of AP#. These numbers were calculated as s_i_ = N δ_i_ p_r_ where δ_i_ is the DS occupancy just before AP# i (Scheuss and Neher, 2001). Of note, there were some variations from one set of experiments to another regarding these parameters, reflecting differences between synapses (**Table 1**). Similar observations have been made at calyx of Held synapses (Lin et al., 2025).

DS and RS replenishment are not independent of each other. A DS can only be filled if an SV is present in the RS, and a downward transition from the RS can only take place if the DS is empty. Likewise, a transition from DS to RS will only happen if the DS is occupied and the RS empty. These considerations are taken into account in the Monte Carlo model. To better understand how the DUs as a whole changed after an AP and during trains, we considered four DU ‘states’ and we followed them throughout the simulations: DUs were “full” when both RS and DS were occupied; “up” when RS was occupied while DS was empty; “down” when RS was empty while DS was occupied; “empty” when both RS and DS were empty (**Supplementary Fig. 3B**). The initial δ and ρ values were used to estimate the initial DU states, keeping in mind the following relations: 1) δ = %down + %full, 2) ρ = %up +%full, and 3) %down + %up + %full + %empty = 1. The DU state was tracked alongside δ and ρ for each simulated AP.

The first AP was then simulated by releasing from the DS that were occupied (DU states full and down). Release resulted in an increase of % up and a simultaneous fall of % full (**Supplementary Fig. 3B,** grey shade).

Below 10 ms, rate constants are largely determined by the large amplitude, fast decaying Ca^2+^ signal occurring near voltage-dependent Ca^2+^ channels (‘Ca^2+^ nanodomain’). The first 10 ms after an AP were simulated with short time intervals (50 μs) while following ρ, δ and the DU states. This timeframe only takes into account forward transitions (r_f_ and s_f_), which were determined from fitting the 100 Hz control data of each dataset, as described before (Miki et al., 2016; Silva et al., 2024). This simulation was then used to establish the overall DU changes in the first 10 ms following an AP. For example, EE was defined as the probability that a DU that was empty just after release would be again empty 10 ms after the AP; EU the probability that a DU that was empty just after release would be up 10 ms after the AP, and so forth (**Table 1**). These probabilities were used to simulate the initial 10 ms of SV replenishment during an ISI for low frequency train simulations. The first 10 ms after release are dominated by the calcium-dependent movement of SVs from the RS to the DS (r_f_, ‘transient docking’) which results in an increase in % down and a decrease in % up (see **Supplementary Fig. 3B**).

After 10 ms, the rate constants depend on the low amplitude, slowly decaying Ca^2+^ signals averaged over the entire varicosity (‘volume-averaged Ca^2+^’). To calculate DU state changes at times > 10 ms after the AP, time intervals for the calculations were changed from 50 μs to 10 ms. This change of time interval was justified because SV movements slow down markedly as a function of time since the AP. It resulted in a marked reduction in calculation time. The transition between the two time domains occurring at 10 ms explains the slight differences between data and simulation points at 10 vs 20 ms in **Fig. 3B**. Whereas transition probabilities were assumed to remain constant during the first 10 ms, they were assumed to change for each subsequent 10 ms time interval as specified in **Supplementary Figure 3A**. In this part of the simulation all transitions were considered, including the back transitions s_b_ and r_b_. Note that the rate r_b_ has a delay before the start of the time-dependent exponential curve and that s_f_ remains constant throughout the ISI. Between 10 and 100 ms after the AP, up and down state proportions were stable, with a low % empty and up DUs and a high % down and full DUs (**Supplementary Figure 3B**; note time scale change at 10 ms). Next, SVs moved from DS to RS (undocking, with rate r_b_), resulting in a gradual increase in % up and a decrease in % down (**Fig. 3B**). As described in the main text, this movement causes a temporary block of replenishment as ρ is high (RS gate closed). DUs gradually returned to their basal state on a time scale on the order of 10 s.

To model DU states during AP trains at low frequency, high time resolution modeling at times 0-10 ms after each AP (time domain 1) was followed by low time resolution modeling starting at 10 ms after the AP and ending at the end of each ISI (time domain 2). Models for consecutive APs were concatenated. For each binding site, the simulation started by attributing a site status to DS and RS. This was done at the beginning of each simulation trial by obtaining a pseudo-random value from the binomial distribution with a mean probability equal to the starting value of δ or ρ, for DS and RS respectively. DU states, δ and ρ were then followed step by step during the entire AP train.

**Supplementary Fig. 3C** depicts the changes in DU state during the simulations of high and low frequency 8-AP trains (**Fig. 2** and **Fig. 4**). The proportion of DUs in each of the full/up/down/empty states is shown alongside ρ and δ just before each AP. During high frequency trains (**left**) an increase in % down (purple continuous curve) at the expense of a decrease in % up (green continuous curve) and % full (orange continuous curve) drives the rise in δ. During low frequency trains (**center**) the trend is the opposite, with undocking causing an increase in the % up (green), mirrored by a reduction in δ. This trend is mainly driven by a drop of % full (orange). DU changes during recovery trains (**right**) reproduce those obtained during control (**left**).

The kinetic parameters were adjusted by hand to find a good fit and to achieve a gradual return to the basal state for long ISIs. The simulation parameter values for the second domain were optimized according to the average release observed in the LFD trains as well as to the awakening train with the final δ and ρ values of the LFD train.

The steady state values of the transition rates in Fig. 3A are heavily constrained by the data within the RS/DS model. By contrast, the time dependent components of r_f_, r_b_ and s_b_, reflecting Ca^2+^- and time-dependent SV movements, are less strictly constrained, due to the limited set of experimental data on the dependence of LFD on ISI.

The states and occupancies were stored for each simulation trial and averages were obtained at the end of the simulation. For the simulations shown, calculations were done 10,000 times for each ISI and resulting averaged values are displayed.

Our LFD simulations reached steady state after a few APs. They are a good model for the first phase of LFD, but they do not account for the second phase of LFD. Modelling the slow phase of LFD would presumably involve changes in rate constants of the model as a function of AP number; such models fall outside the scope of the present work.

### Failure analysis

For this analysis we focused on the first component of LFD and more specifically, on the value of s_2_. Results were taken from experiments with ISIs of 500 ms (**Fig. 2**), 800 ms and 1.6 s (**Fig. 1**). The trains were sorted according to whether the 1st AP failed or was successful (**Fig. 5A**; failures in purple, successes in black). The ratio of the released SV number after AP# 2 (s_2_) and the average of the total released SV number after AP# 1 (<s_1_>) was calculated for the s_1_ failures and for the s_1_ successes separately. One synapse had no failures; it was excluded from this analysis. Both traces with successes and failures show depression, as evidenced by a s_2_/<s_1_> (PPR) lower than 1, indicating that loss of SVs is not the main mechanism of LFD (**Fig. 5B, left**).

This analysis was extended by sorting the trains according to the number of SVs released for s_1_. Though the data shows a trend of deeper depression with increased release during the 1st AP, the difference is not significant (**Fig. 5B, right**), indicating again that while release may play a role it is not the main mechanism. Next, we evaluated if the simulations replicated this characteristic of the data. The simulation trials were sorted the same way as the data and the PPRs calculated for each case (**Fig. 5B**; see black dots). The simulation also shows a significant depression for successful trials, with a somewhat stronger correlation between the amount of depression and the initial s_1_ value than experimentally observed. Overall, the simulation reproduces well the results of the experiments.

### Binomial analysis and predictions

According to the binomial distribution, the probability that s_i_, the number of SVs released after AP number i, equals k is p(k) = P_i_^k^ (1 − P_i_)^N−k^ N!/(k!(N − k)!), where P_i_ is the release probability per docking site (P_i_ = δ_i_ p_r_) and N is the number of docking sites (Malagon et al., 2016). The values of N and P were obtained by minimizing the summed squared deviations between the binomial prediction and data. This fitting was carried out for the control s_1_ data of the synapses in **Fig 7**, giving the distribution p_ctrl_(k). LFD was evaluated in a similar manner, considering each of the stimuli in the train as a separate event. The distributions of released SV numbers for 2 examples of control s_1_ data is shown in black lines in **Supplementary Fig. 5A** and red lines show the LFD data distributions, p_LFD_(k).

We tested the case of the depression in LFD being the result of an increase in stimulus failures, with a constant probability of release. In this case the probability of failures during LFD, p_LFD_(0), would be larger than the control probability of failures, p_ctl_(0), but the rest of the distribution would retain the same shape for the LFD and control distributions: p_LFD_(k) / p_ctl_(k) = constant for k = 1 to N.

Considering that for each distribution, the sum of p(k) from 0 to N is 1, we obtain: p_LFD_(k) = p_ctl_(k) x [1-( p_LFD_(0)- p_ctl_(0))/(1- p_ctl_(0))] for k = 1 to N.

The dotted blue line in the examples of **Supplementary Fig. 5A** show the result of this prediction.

If, instead, LFD was the result of a decrease in probability of release then the p(k) of the LFD data would follow the distribution: p_LFD_(k) = (P_LFD_)^k^ x (1 - P_LFD_)^N-k^ N!/(k!(N − k)!)

where N is the number of DS (assumed to be 5 in the example shown) and P_LFD_ is the P calculated from the LFD distribution: P_LFD_ = 1 - p_LFD_(0)^1/N^. The dotted purple lines in the examples of **Supplementary Fig. 5A** show the result of this prediction. The LFD data (red line) was then compared to both predictions. In both examples the decreased P prediction (purple) is much closer to the data than the prediction for increased stimulus failure (blue). To quantify this difference residuals were calculated for each point in the plot for both predictions. The group data for the residuals of each prediction is shown in **Supplementary Fig. 5B, left**. The right panel depicts the average of the residuals. The prediction of a decrease in P is much closer to the data, compared to the prediction for an increase in the number of stimulus failures.

This analysis shows that an increase in stimulus failures does not account for the distribution of events in the LFD trials and that a decrease in P explains this phenomenon better.

## Abbreviations

AP: Action Potential
AZ: Active Zone
DS: Docking Site
DU: Docking Unit
HFD: High Frequency Depression
IP: Intermediate Pool
ISI: Inter-Stimulus Interval
LFD: Low Frequency Depression
MLI: Molecular Layer Interneuron
PF: Parallel Fiber
PPR: Paired-Pulse Ratio
RRP: Readily Releasable Pool
RS: Replacement Site
STP: Short Term Synaptic Plasticity
SV: Synaptic Vesicle
δ: Docking site occupancy
ρ: Replacement site occupancy

## Acknowledgments

We thank Takafumi Miki and Shin-ya Kawaguchi for their help in the modelling. This work was supported by CNRS (UMR 8003), by the European Research Council (Advanced Grant SinSyn to A. M.), and by Agence Nationale pour la Recherche (Grant PLASTICAZ to A. M.).

## Supplementary Material

### Supplementary figures Legends

**Supplementary Figure 1:**
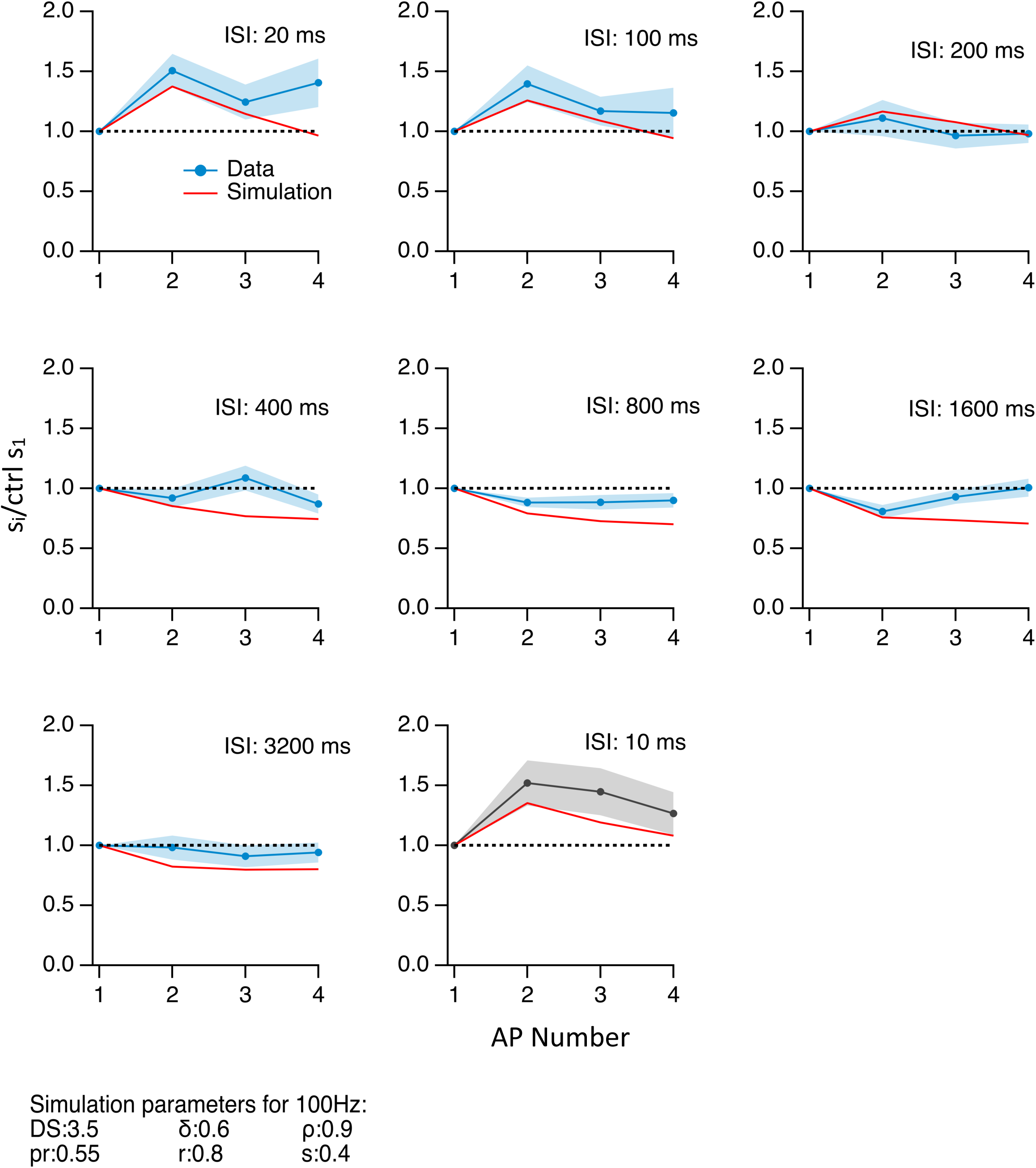
Trains of 4 successive EPSCs with different ISIs. Each panel shows numbers of released SVs in response to 4 consecutive APs at different ISI values. These numbers are normalized with respect to the mean s_1_ value, calculated across all ISI values (dotted lines). Data (dots: mean values; shaded areas: ± sem) are in blue for ISI values of 20 to 3200 ms, and in black for control 8-AP trains with an ISI of 10 ms. Red curves show simulations based on the model of Fig. 3.

**Supplementary Figure 2:**
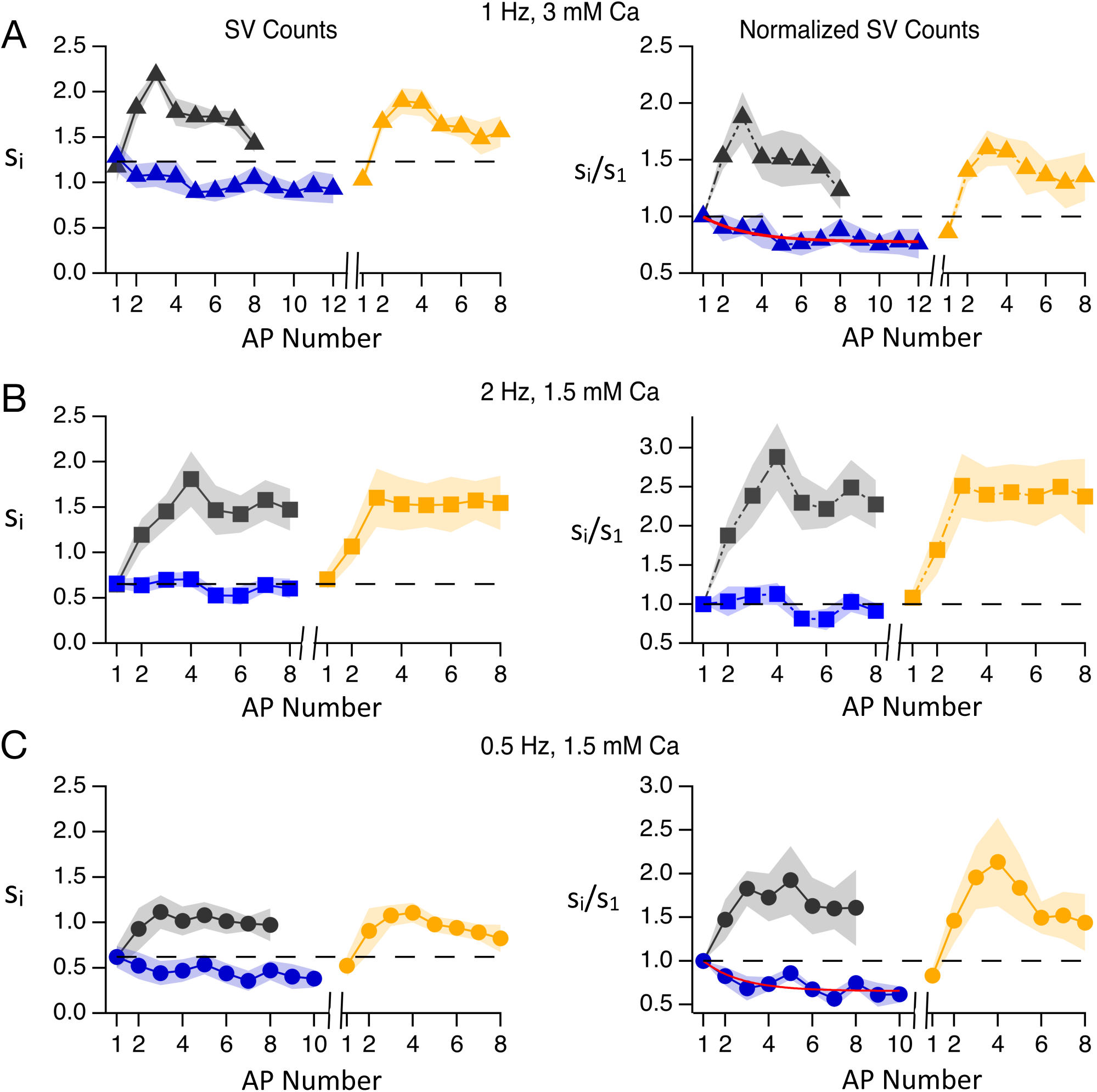
LFD at various ISI and external calcium concentration values. Same experiments as in Fig. 2 at varying stimulation frequencies during low frequency trains (**A**: 1 Hz, n = 5; **B**: 2 Hz, n = 6; **C**: 0.5 Hz, n = 6) and varying external calcium concentrations (**A**: 3 mM; **B** and **C**: 1.5 mM). Red curves show exponential fits (**A**: asymptote: 0.79 ± 0.08; τ = 1.92 ± 0.55 ISIs, or 1.92 s; C: asymptote: 0.69; τ = 1.8 ISIs, or 3.6 s).

**Supplementary Figure 3:**
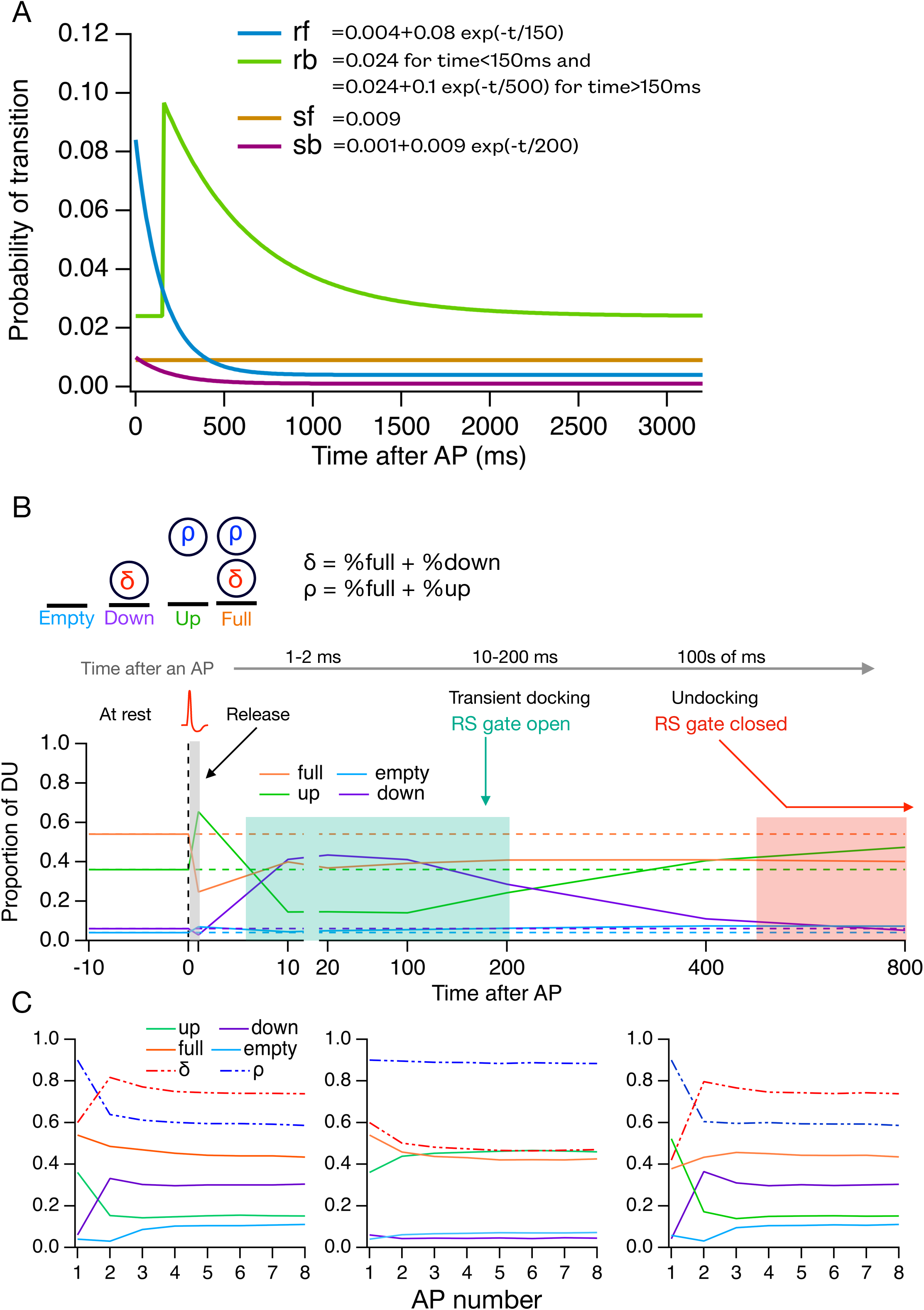
Time dependent transition rates during LFD. **A**: Time dependent transition rates (calculated per 10 ms time bins) are shown for the 4 SV transitions of the model in Fig. 2A. Note that the first 10 ms period is excluded from this plot (see the corresponding transition probabilities in **Table 1**). s_f_ is stable as a function of time, while r_f_, r_b_ and s_b_ follow exponential functions of time (t, in ms). In the case of r_b_, the exponential starts with a delay of 150 ms. **B**: Following an AP, SV movements were described using 4 different DU states: “full” when both RS and DS were occupied; “up” when RS was occupied while DS was empty; “down” when RS was empty while DS was occupied; “empty” when both RS and DS were empty. Release resulted in an increase of % up and a simultaneous fall of % full (gray shade). The next 10 ms following stimulation were dominated by the calcium-dependent movement of SVs from the RS to the DS (r_f_, ‘transient docking’) resulting in an increase in % down and a decrease in % up. In subsequent 10 ms time segments, SV movements were calculated based on the time-dependent changes in r_f_, r_b_, s_f_ and s_b_ shown in **A**. Between 10 and 100 ms after the AP, up and down state proportions were stable, with a low % empty and up DUs and a high % down and full DUs (green shade, RS gate open). Next, SVs moved from DS to RS (undocking, with rate r_b_), resulting in a gradual increase in % up and a decrease in % down (red shade, RS gate closed). **C**: DU states before individual APs for 8-AP trains. **Left**: Control. **Center**: LFD 2 Hz. **Right**: Recovery.

**Supplementary Figure 4:**
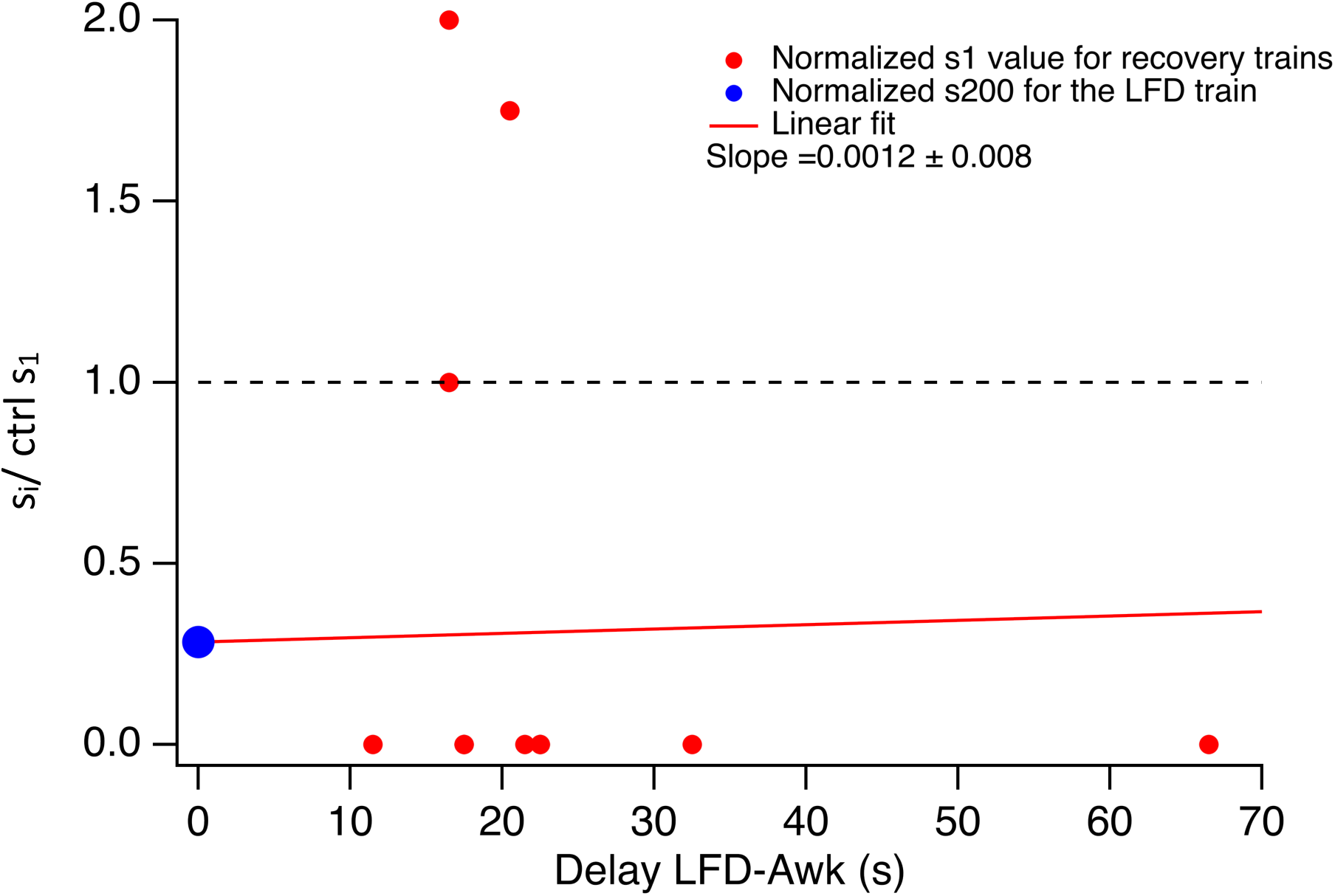
Lack of recovery of LFD over tens of s after AP train. The time interval between the last AP in the LFD train and the 1st AP in the awakening train varied in the experiments presented in Fig. 7. These differences were used to analyze the time it takes for synapses to recover after a long low-frequency train. This figure shows the normalized s_1_ value for recovery trains following long LFD trains (200 APs @ 2 Hz), as a function of the time duration intervening between the end of the LFD train and the onset of the recovery train. Failures were observed in a majority of the trials. The 0-delay point (blue dot) corresponds to the mean s_200_ value at the end of the LFD train. The red line is a linear fit of the data that is constrained to pass through the 0-delay point. The data indicate no recovery for up to 60 s.

**Supplementary Figure 5:**
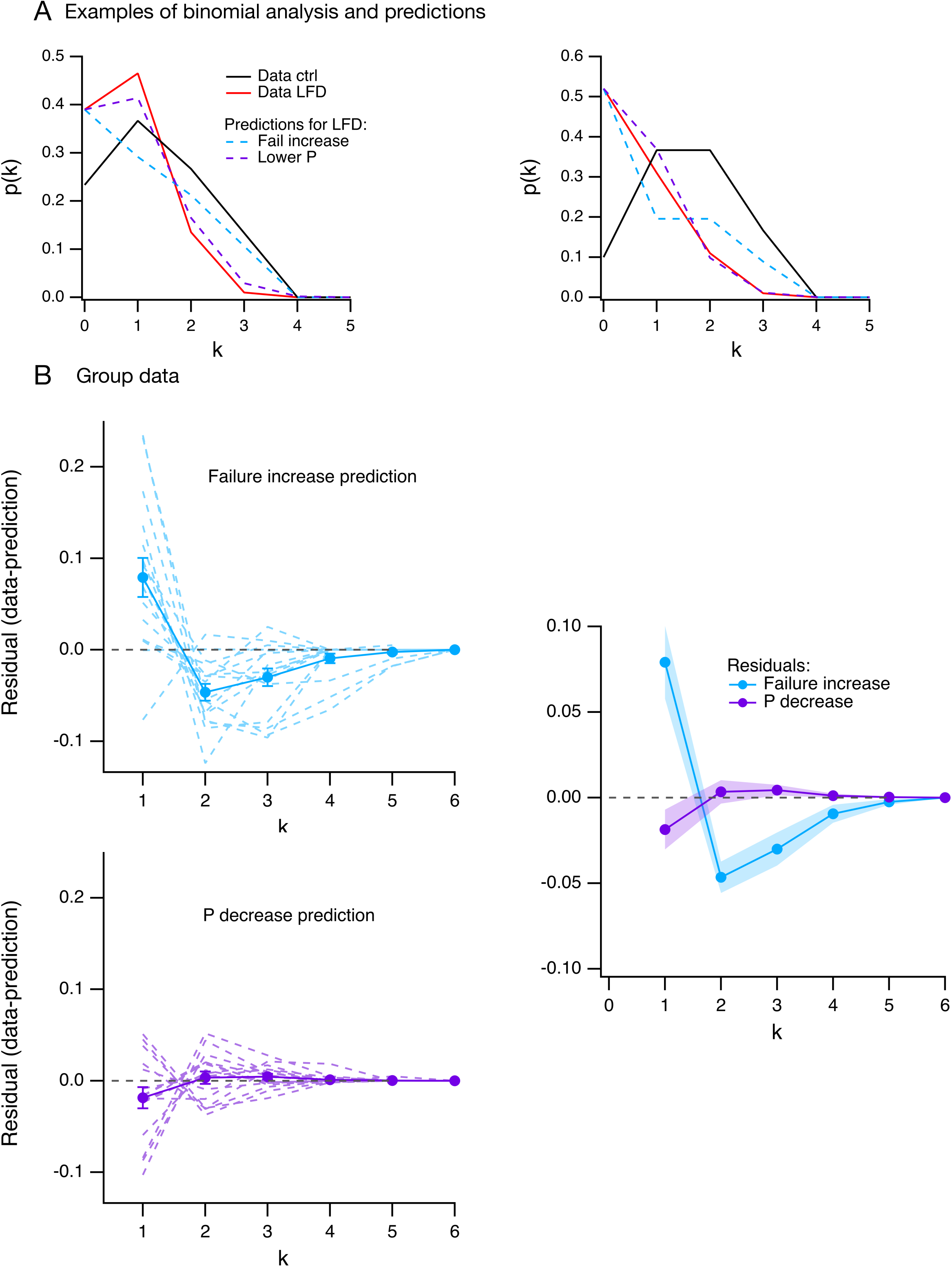
Binomial analysis of released SV numbers during prolonged LFD. **A**: Exemplar experiments. In each case, the distribution of released SV numbers (k) in response to the 1^st^ AP during control runs (8 APs @ 100 Hz; black curves) is shown together with the distribution of released SV numbers during LFD (red curves; obtained from two of the experiments shown in Fig. 7). Next, we calculated the binomial distributions in two different models of reduced responsiveness. Both models assumed unchanged N values. In one model (blue dotted curves), the reduced response was assumed to reflect global failure at the AZ level for some stimulations, while other stimulations were assumed to be as effective as in the control. This model mimics situations such as unwanted stimulation failures during LFD. In the other model, a homogeneous reduction of P (= δ* p_r_; the release probability per DU) was assumed (purple dotted curves). Both models were constrained to account for the failure rate observed during LFD. In both experiments, the lower P curve is much closer to the experimental results (red curves for k = 1 to 3) than the failure increase curve. **B**: Group data analysis (n = 16 trials from 8 cells). **Left**: Residuals (dotted lines: individual LFD trials; dots and associated SEM, means across trials) calculated from the comparison of the two models with LFD data (**top:** increased failure model; **bottom:** decreased P model). **Right**: Superimposition of mean residual curves for the two models, showing that the lower P model is a much better representation of the data than the failure model.

**Supplementary Figure 6:**
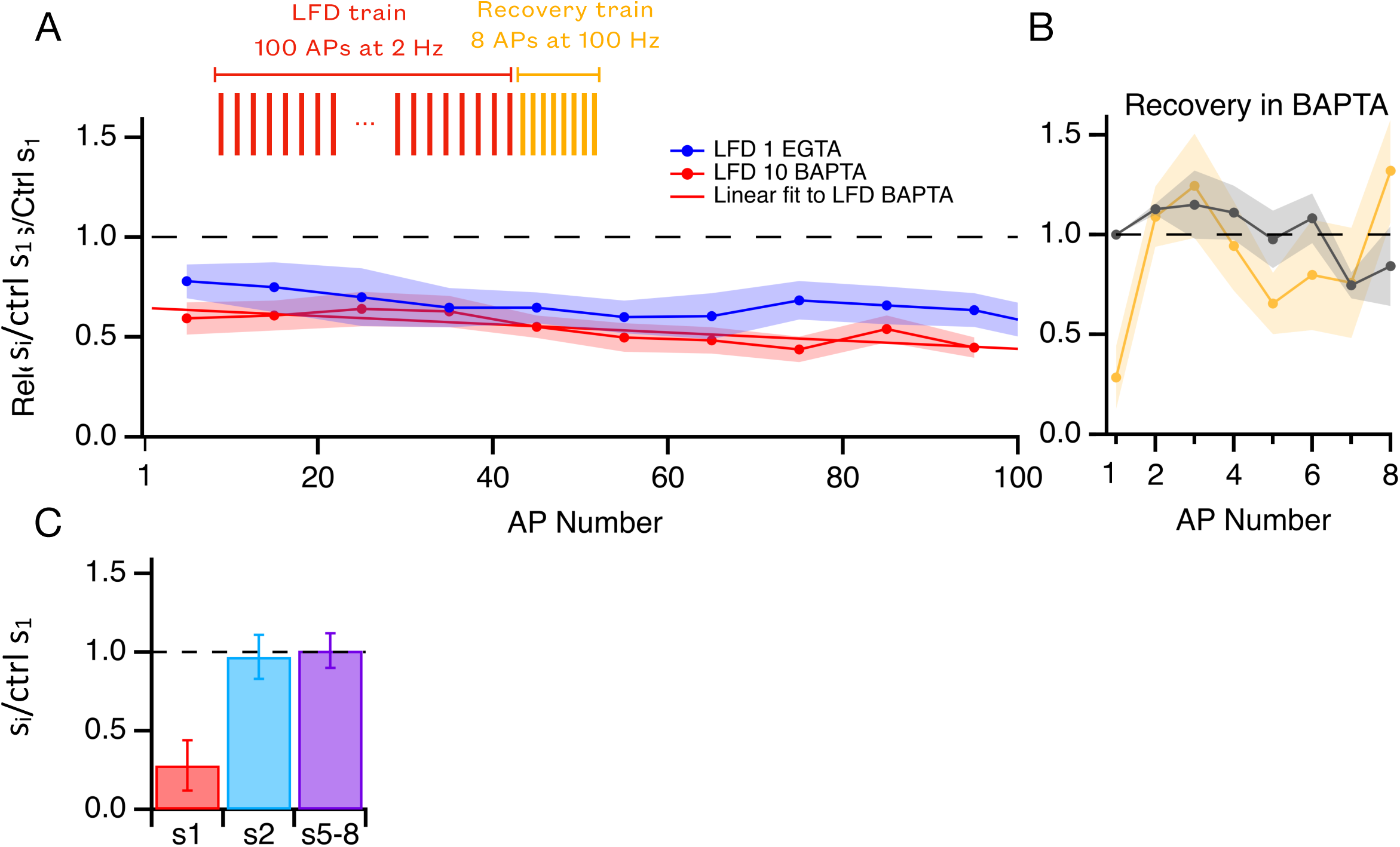
Prolonged low frequency trains with postsynaptic BAPTA. BAPTA (10 mM) was added to the internal solution used for MLI recording, keeping all other experimental conditions as before. **A:** A long AP train at low frequency (100 APs @ 2 Hz) was followed by a short 8-AP recovery train (experimental protocol in **insert**). Normalized plot of s_i_ during long low frequency train (red curve showing binned data for 10 consecutive APs), showing a two-component depression with an initial rapid depression (65 ± 3 % of the control) followed by a gradual depression during the entire train duration (slope = -0.2 ± 0.05 % per AP; red: linear fit to the data ; n = 12 trials, 3 cells). The results resemble those with 1 mM EGTA shown in Fig. 6 (reproduced here as a dark blue curve). **B**: A short (8-AP) 100 Hz recovery train was applied immediately at the end of the low-frequency train. Normalized s_i_ plots for control and the recovery 100 Hz trains are compared here. **C:** The s_1_ ratio between recovery and control trains is < 1, indicating a decrease in δ at the end of the low frequency train while neither s_2_ (blue) nor s_5-8_ (purple) are changed, suggesting no change in ρ or in IP size. The similarity of results with and without postsynaptic BAPTA supports a presynaptic mechanism for LFD.

## References

Abrahamsson T, Gustafsson B, Hanse E. 2007. Reversible synaptic depression in developing rat CA3–CA1 synapses explained by a novel cycle of AMPA silencing-unsilencing. Journal of Neurophysiology 98:2604–2611. DOI: 10.1152/jn.00602.2007, PMID: 17804578

Atluri PP, Regehr WG. 1998. Delayed release of neurotransmitter from cerebellar granule cells. The Journal of Neuroscience 18:8214–8227. DOI: 10.1523/jneurosci.18-20-08214.1998

Betz WJ. 1970. Depression of transmitter release at the neuromuscular junction of the frog. The Journal of Physiology 206:629–644. DOI: 10.1113/jphysiol.1970.sp009034, PMID: 5498509

Blanchard K, Martín JZ de S, Marty A, Llano I, Trigo FF. 2020. Differentially poised vesicles underlie fast and slow components of release at single synapses. Journal of General Physiology 152:e201912523. DOI: 10.1085/jgp.201912523, PMID: 32243497

Bolshakov AP, Kolleker A, Volkova EP, Valiullina-Rakhmatullina F, Kolosov PM, Rozov A. 2019. Overexpression of calretinin enhances short-term synaptic depression. Frontiers in Cellular Neuroscience 13:91. DOI: 10.3389/fncel.2019.00091, PMID: 30930749

Borges-Merjane C, Kim O, Jonas P. 2020. Functional electron microscopy, “Flash and Freeze,” of identified cortical synapses in acute brain slices. Neuron 105:992–1006.e6. DOI: 10.1016/j.neuron.2019.12.022, PMID: 31928842

Brenowitz SD, Regehr WG. 2007. Reliability and heterogeneity of calcium signaling at single presynaptic boutons of cerebellar granule cells. The Journal of Neuroscience 27:7888–7898. DOI: 10.1523/jneurosci.1064-07.2007, PMID: 17652580

Buzsáki G, Mizuseki K. 2014. The log-dynamic brain: how skewed distributions affect network operations. Nature Reviews Neuroscience 15:264–278. DOI: 10.1038/nrn3687, PMID: 24569488

Casado M, Dieudonné S, Ascher P. 2000. Presynaptic N-methyl-d-aspartate receptors at the parallel fiber–Purkinje cell synapse. Proceedings of the National Academy of Sciences 97:11593–11597. DOI: 10.1073/pnas.200354297, PMID: 11016958

Castillo J del, Katz B. 1954. Statistical factors involved in neuromuscular facilitation and depression. The Journal of Physiology 124:574–585. DOI: 10.1113/jphysiol.1954.sp005130, PMID: 13175200

Chadderton P, Margrie TW, Häusser M. 2004. Integration of quanta in cerebellar granule cells during sensory processing. Nature 428:856–860. DOI: 10.1038/nature02442, PMID: 15103377

Charlton MP, Smith SJ, Zucker RS. 1982. Role of presynaptic calcium ions and channels in synaptic facilitation and depression at the squid giant synapse. The Journal of Physiology 323:173–193. DOI: 10.1113/jphysiol.1982.sp014067, PMID: 6284915

Chiu DN, Carter BC. 2024. Synaptotagmin 7 sculpts short-term plasticity at a high probability synapse. The Journal of Neuroscience 44:e1756232023. DOI: 10.1523/jneurosci.1756-23.2023, PMID: 38262726

Dittman JS, Kreitzer AC, Regehr WG. 2000. Interplay between facilitation, depression, and residual calcium at three presynaptic terminals. The Journal of Neuroscience 20:1374–1385. DOI: 10.1523/jneurosci.20-04-01374.2000

Doussau F, Humeau Y, Benfenati F, Poulain B. 2010. A novel form of presynaptic plasticity based on the fast reactivation of release sites switched off during low-frequency depression. The Journal of Neuroscience 30:16679–16691. DOI: 10.1523/jneurosci.3644-09.2010, PMID: 21148007

Doussau F, Schmidt H, Dorgans K, Valera AM, Poulain B, Isope P. 2017. Frequency-dependent mobilization of heterogeneous pools of synaptic vesicles shapes presynaptic plasticity. eLife 6:e28935. DOI: 10.7554/elife.28935, PMID: 28990927

Eshra A, Schmidt H, Eilers J, Hallermann S. 2021. Calcium dependence of neurotransmitter release at a high fidelity synapse. eLife 10:e70408. DOI: 10.7554/elife.70408, PMID: 34612812

Fukaya, R., Hirai, H., Sakamoto, H., Hashimotodani, Y., Hirose, K., and Sakaba, T. (2023). Increased vesicle fusion competence underlies long-term potentiation at hippocampal mossy fiber synapses. Sci. Adv. 9, eadd3616. 10.1126/sciadv.add3616.

Huson V, Regehr WG. 2020. Diverse roles of Synaptotagmin-7 in regulating vesicle fusion. Current Opinion in Neurobiology 63:42–52. DOI: 10.1016/j.conb.2020.02.006, PMID: 32278209

Imig C, Min S-W, Krinner S, Arancillo M, Rosenmund C, Südhof TC, Rhee J, Brose N, Cooper BH. 2014. The morphological and molecular nature of synaptic vesicle priming at presynaptic active zones. Neuron 84:416--431. DOI: 10.1016/j.neuron.2014.10.009, PMID: 25374362

Jackman SL, Turecek J, Belinsky JE, Regehr WG. 2016. The calcium sensor synaptotagmin 7 is required for synaptic facilitation. Nature 529:88–91. DOI: 10.1038/nature16507, PMID: 26738595

Jörntell H, Ekerot C-F. 2003. Receptive field plasticity profoundly alters the cutaneous parallel fiber synaptic input to cerebellar interneurons in vivo. The Journal of Neuroscience 23:9620–9631. DOI: 10.1523/jneurosci.23-29-09620.2003

Kavalali ET. 2006. Synaptic vesicle reuse and its implications. The Neuroscientist 12:57–66. DOI: 10.1177/1073858405281852, PMID: 16394193

Kim O, Okamoto Y, Kaufmann WA, Brose N, Shigemoto R, Jonas P. 2024. Presynaptic cAMPPKA-mediated potentiation induces reconfiguration of synaptic vesicle pools and channel-vesicle coupling at hippocampal mossy fiber boutons. PLOS Biology 22:e3002879. DOI: 10.1371/journal.pbio.3002879, PMID: 39556620

Kobbersmed JRL, Grasskamp AT, Jusyte M, Böhme MA, Ditlevsen S, Sørensen JB, Walter AM. 2020. Rapid regulation of vesicle priming explains synaptic facilitation despite heterogeneous vesicle:Ca2+ channel distances. eLife 9:e51032. DOI: 10.7554/elife.51032, PMID: 32077852

Koppensteiner P, Bhandari P, Önal C, Borges-Merjane C, Monnier EL, Nakamura Y, Sadakata T, Sanbo M, Hirabayashi M, Brose N, Jonas P, Shigemoto R. 2022. A two-pool mechanism of vesicle release in medial habenula terminals underlies GABAB receptor-mediated potentiation. bioRxiv 2022.10.28.514202. DOI: 10.1101/2022.10.28.514202

Kusick GF, Chin M, Raychaudhuri S, Lippmann K, Adula KP, Hujber EJ, Vu T, Davis MW, Jorgensen EM, Watanabe S. 2020. Synaptic vesicles transiently dock to refill release sites. Nature Neuroscience 23:1329–1338. DOI: 10.1038/s41593-020-00716-1, PMID: 32989294

Kusick GF, Ogunmowo TH, Watanabe S. 2022. Transient docking of synaptic vesicles: Implications and mechanisms. Current Opinion in Neurobiology 74:102535. DOI: 10.1016/j.conb.2022.102535, PMID: 35398664

Lin K, Ranjan M, Lipstein N, Brose N, Neher E, Taschenberger H. 2025. Number and relative abundance of synaptic vesicles in functionally distinct priming states determine synaptic strength and short-term plasticity. The Journal of Physiology. DOI: 10.1113/jp286282, PMID: 40120134

Lin K-H, Taschenberger H, Neher E. 2022. A sequential two-step priming scheme reproduces diversity in synaptic strength and short-term plasticity. Proceedings of the National Academy of Sciences 119:e2207987119. DOI: 10.1073/pnas.2207987119, PMID: 35969787

MacDougall DD, Lin Z, Chon NL, Jackman SL, Lin H, Knight JD, Anantharam A. 2018. The highaffinity calcium sensor synaptotagmin-7 serves multiple roles in regulated exocytosis. Journal of General Physiology 150:783–807. DOI: 10.1085/jgp.201711944, PMID: 29794152

Malagon G, Miki T, Llano I, Neher E, Marty A. 2016. Counting Vesicular Release Events Reveals Binomial Release Statistics at Single Glutamatergic Synapses. The Journal of Neuroscience 36:4010–4025. DOI: 10.1523/jneurosci.4352-15.2016, PMID: 27053208

Malagon G, Miki T, Tran V, Gomez LC, Marty A. 2020. Incomplete vesicular docking limits synaptic strength under high release probability conditions. eLife 9:e52137. DOI: 10.7554/elife.52137, PMID: 32228859

Margrie TW, Brecht M, Sakmann B. 2002. In vivo, low-resistance, whole-cell recordings from neurons in the anaesthetized and awake mammalian brain. Pflügers Archiv 444:491–498. DOI: 10.1007/s00424-002-0831-z, PMID: 12136268

Miki T, Kaufmann WA, Malagon G, Gomez L, Tabuchi K, Watanabe M, Shigemoto R, Marty A. 2017. Numbers of presynaptic Ca 2+ channel clusters match those of functionally defined vesicular docking sites in single central synapses. Proceedings of the National Academy of Sciences 114:E5246–E5255. DOI: 10.1073/pnas.1704470114, PMID: 28607047

Miki T, Malagon G, Pulido C, Llano I, Neher E, Marty A. 2016. Actin- and myosin-dependent vesicle loading of presynaptic docking sites prior to exocytosis. Neuron 91:808–823. DOI: 10.1016/j.neuron.2016.07.033, PMID: 27537485

Miki T, Midorikawa M, Sakaba T. 2020. Direct imaging of rapid tethering of synaptic vesicles accompanying exocytosis at a fast central synapse. Proceedings of the National Academy of Sciences 117:14493–14502. DOI: 10.1073/pnas.2000265117, PMID: 32513685

Miki T, Nakamura Y, Malagon G, Neher E, Marty A. 2018. Two-component latency distributions indicate two-step vesicular release at simple glutamatergic synapses. Nature Communications 9:3943. DOI: 10.1038/s41467-018-06336-5, PMID: 30258069

Müller M, Goutman JD, Kochubey O, Schneggenburger R. 2010. Interaction between Facilitation and Depression at a Large CNS Synapse Reveals Mechanisms of Short-Term Plasticity. The Journal of Neuroscience 30:2007–2016. DOI: 10.1523/jneurosci.4378-09.2010, PMID: 20147529

Neher E. 2023. Interpretation of presynaptic phenotypes of synaptic plasticity in terms of a two-step priming process. Journal of General Physiology 156:e202313454. DOI: 10.1085/jgp.202313454, PMID: 38112713

Neher E. 2015. Merits and limitations of vesicle pool models in view of heterogeneous populations of synaptic vesicles. Neuron 87:1131–1142. DOI: 10.1016/j.neuron.2015.08.038, PMID: 26402599

Neher E, Brose N. 2018. Dynamically primed synaptic vesicle states: Key to understand synaptic short-term plasticity. Neuron 100:1283–1291. DOI: 10.1016/j.neuron.2018.11.024, PMID: 30571941

Neher E, Sakaba T. 2008. Multiple roles of calcium ions in the regulation of neurotransmitter release. Neuron 59:861–872. DOI: 10.1016/j.neuron.2008.08.019, PMID: 18817727

Ogunmowo TH, Hoffmann C, Patel C, Pepper R, Wang H, Gowrisankaran S, Idel J, Ho A, Raychaudhuri S, Maher BJ, Cooper BH, Milosevic I, Milovanovic D, Watanabe S. 2025. Intersectin and endophilin condensates prime synaptic vesicles for release site replenishment. Nature Neuroscience 28:1649–1662. DOI: 10.1038/s41593-025-02002-4, PMID: 40629141

Pulido C, Marty A. 2018. A two-step docking site model predicting different short-term synaptic plasticity patterns. Journal of General Physiology 150:1107–1124. DOI: 10.1085/jgp.201812072, PMID: 29950400

Rebola N, Reva M, Kirizs T, Szoboszlay M, Lőrincz A, Moneron G, Nusser Z, DiGregorio DA. 2019. Distinct nanoscale calcium channel and synaptic vesicle topographies contribute to the diversity of synaptic function. Neuron 104:693–710.e9. DOI: 10.1016/j.neuron.2019.08.014, PMID: 31558350

Redman RS, Silinsky EM. 1994. ATP released together with acetylcholine as the mediator of neuromuscular depression at frog motor nerve endings. The Journal of Physiology 477:117–127. DOI: 10.1113/jphysiol.1994.sp020176, PMID: 8071878

Regehr WG. 2012. Short-term presynaptic plasticity. Cold Spring Harbor Perspectives in Biology 4:a005702. DOI: 10.1101/cshperspect.a005702, PMID: 22751149

Ritzau-Jost A, Delvendahl I, Rings A, Byczkowicz N, Harada H, Shigemoto R, Hirrlinger J, Eilers J, Hallermann S. 2014. Ultrafast action potentials mediate kilohertz signaling at a central synapse. Neuron 84:152–163. DOI: 10.1016/j.neuron.2014.08.036, PMID: 25220814

Ritzau-Jost A, Jablonski L, Viotti J, Lipstein N, Eilers J, Hallermann S. 2018. Apparent calcium dependence of vesicle recruitment. The Journal of Physiology 596:4693–4707. DOI: 10.1113/jp275911, PMID: 29928766

Rossi DJ, Alford S, Mugnaini E, Slater NT. 1995. Properties of transmission at a giant glutamatergic synapse in cerebellum: the mossy fiber-unipolar brush cell synapse. Journal of Neurophysiology 74:24–42. DOI: 10.1152/jn.1995.74.1.24, PMID: 7472327

Rudolph S, Overstreet-Wadiche L, Wadiche JI. 2011. Desynchronization of multivesicular release enhances Purkinje cell output. Neuron 70:991–1004. DOI: 10.1016/j.neuron.2011.03.029, PMID: 21658590

Saviane C, Savtchenko LP, Raffaelli G, Voronin LL, Cherubini E. 2002. Frequency-dependent shift from paired-pulse facilitation to paired-pulse depression at unitary CA3-CA3 synapses in the rat hippocampus. The Journal of Physiology 544:469–476. DOI: 10.1113/jphysiol.2002.026609, PMID: 12381819

Scheuss V, Neher E. 2001. Estimating synaptic parameters from mean, variance, and covariance in trains of synaptic responses. Biophysical Journal 81:1970–1989. DOI: 10.1016/s0006-3495(01)75848-1, PMID: 11566771

Schmidt H. 2019. Control of presynaptic parallel fiber efficacy by activity-dependent regulation of the number of occupied release sites. Frontiers in Systems Neuroscience 13:30. DOI: 10.3389/fnsys.2019.00030, PMID: 31379524

Siksou L, Varoqueaux F, Pascual O, Triller A, Brose N, Marty S. 2009. A common molecular basis for membrane docking and functional priming of synaptic vesicles. European Journal of Neuroscience 30:49–56. DOI: 10.1111/j.1460-9568.2009.06811.x, PMID: 19558619

Silva M, Tran V, Marty A. 2024. A maximum of two readily releasable vesicles per docking site at a cerebellar single active zone synapse. eLife 12:RP91087. DOI: 10.7554/elife.91087, PMID: 38180320

Silva M, Tran V, Marty A. 2021. Calcium-dependent docking of synaptic vesicles. Trends in Neurosciences 44:579–592. DOI: 10.1016/j.tins.2021.04.003, PMID: 34049722

Silverman-Gavrila LB, Orth PMR, Charlton MP. 2005. Phosphorylation-dependent lowfrequency depression at phasic synapses of a crayfish motoneuron. The Journal of Neuroscience 25:3168–3180. DOI: 10.1523/jneurosci.4908-04.2005, PMID: 15788774

Soler-Llavina GJ, Sabatini BL. 2006. Synapse-specific plasticity and compartmentalized signaling in cerebellar stellate cells. Nature Neuroscience 9:798–806. DOI: 10.1038/nn1698, PMID: 16680164

Tan YP, Llano I, Hopt A, Würriehausen F, Neher E. 1999. Fast scanning and efficient photodetection in a simple two-photon microscope. Journal of Neuroscience Methods 92:123–135. DOI: 10.1016/s0165-0270(99)00103-x, PMID: 10595710

Tanaka M, Sakaba T, Miki T. 2021. Quantal analysis estimates docking site occupancy determining short-term depression at hippocampal glutamatergic synapses. The Journal of Physiology 599:5301–5327. DOI: 10.1113/jp282235, PMID: 34705277

Taschenberger H, Woehler A, Neher E. 2016. Superpriming of synaptic vesicles as a common basis for intersynapse variability and modulation of synaptic strength. Proceedings of the National Academy of Sciences 113:E4548–E4557. DOI: 10.1073/pnas.1606383113, PMID: 27432975

Tran V, Miki T, Marty A. 2022. Three small vesicular pools in sequence govern synaptic response dynamics during action potential trains. Proceedings of the National Academy of Sciences 119:e2114469119. DOI: 10.1073/pnas.2114469119, PMID: 35101920

Tran V, Silva M, Marty A. 2023. Prioritized docking of synaptic vesicles provided by a rapid recycling pathway. iScience 26:106366. DOI: 10.1016/j.isci.2023.106366, PMID: 37009220

Trommershäuser J, Schneggenburger R, Zippelius A, Neher E. 2003. Heterogeneous presynaptic release probabilities: Functional relevance for short-term plasticity. Biophysical Journal 84:1563–1579. DOI: 10.1016/s0006-3495(03)749674, PMID: 12609861

Trussell LO, Zhang S, Raman IM. 1993. Desensitization of AMPA receptors upon multiquantal neurotransmitter release. Neuron 10:1185–1196. DOI: 10.1016/08966273(93)90066-z, PMID: 7686382

Turecek J, Jackman SL, Regehr WG. 2017. Synaptotagmin 7 confers frequency invariance onto specialized depressing synapses. Nature 551:503–506. DOI: 10.1038/nature24474, PMID: 29088700

Turecek J, Jackman SL, Regehr WG. 2016. Synaptic specializations support frequency-independent Purkinje cell output from the cerebellar cortex. Cell Reports 17:3256–3268. DOI: 10.1016/j.celrep.2016.11.081, PMID: 28009294

Weingarten DJ, Shrestha A, Juda-Nelson K, Kissiwaa SA, Spruston E, Jackman SL. 2022. Fast resupply of synaptic vesicles requires synaptotagmin-3. Nature 611:320–325. DOI: 10.1038/s41586-022-05337-1, PMID: 36261524

Wölfel M, Lou X, Schneggenburger R. 2007. A mechanism intrinsic to the vesicle fusion machinery determines fast and slow transmitter release at a large CNS synapse. The Journal of Neuroscience 27:3198–3210. DOI: 10.1523/jneurosci.4471-06.2007, PMID: 17376981

Wu L-G, Borst JGG. 1999. The reduced release probability of releasable vesicles during recovery from short-term synaptic depression. Neuron 23:821–832. DOI: 10.1016/s0896-6273(01)80039-8, PMID: 10482247

Wu Z, Kusick GF, Berns MM, Raychaudhuri S, Itoh K, Walter AM, Chapman ER, Watanabe S. 2024. Synaptotagmin 7 docks synaptic vesicles to support facilitation and Doc2αtriggered asynchronous release. eLife 12:RP90632. DOI: 10.7554/elife.90632, PMID: 38536730

Xu J, Wu L-G. 2005. The decrease in the presynaptic calcium current Is a major cause of short-term depression at a calyx-type synapse. Neuron 46:633–645. DOI: 10.1016/j.neuron.2005.03.024, PMID: 15944131

Zucker RS, Regehr WG. 2002. Short-term synaptic plasticity. Annual Review of Physiology 64:355--405. DOI: 10.1146/annurev.physiol.64.092501.114547, PMID: 11826273

